# Tracking Satb2-positive retinal ganglion cells in zebrafish unveils developmental functional reorganization

**DOI:** 10.64898/2025.12.25.696470

**Authors:** Ayjan Urazbayeva, Fumi Kubo

## Abstract

Retinal ganglion cells (RGCs) relay visual information to the brain, and their functional properties are crucial for visually guided behavior. However, functional maturation of RGCs across development remains elusive. Here, we investigated the developmental maturation of a genetically defined RGC subtype expressing Satb2 in zebrafish, which allowed single-layer-resolved analysis of the tectal neuropil. Using *in vivo* calcium imaging, we characterized visual response properties of Satb2-positive RGC axon terminals at two developmental stages. At the early stage, responses segregated into seven functional clusters with diverse tuning profiles. By later stage, these responses reorganized into five clusters, indicating developmental restructuring of functional identities. Longitudinal imaging of individual RGC axons revealed that while some response types remain stable across development, others exhibit pronounced plasticity. In particular, certain RGC cluster acquired additional tuning features and converged into motion-sensitive cluster, coinciding with a marked expansion of motion-responsive cluster and reflecting progressive functional specialization. Visual deprivation experiments revealed that core visual tuning is genetically determined, whereas visual experience is crucial for fine-tuning of the particular functional subtype. Together, our findings provide a single-cell-resolved view of retinal circuit maturation and reveal how stable and flexible responses are integrated during the development of vertebrate vision.

## Introduction

Visual circuits extract behaviorally relevant features from the external environment through parallel processing that begins in the retina. Retinal ganglion cells (RGCs), the sole output neurons of the retina, transmit information to downstream brain areas, enabling visually guided behaviors. Advances in anatomical characterizations, functional imaging, and transcriptomics have revealed that RGCs comprise a large number of distinct subtypes, each characterized by specific morphological features, visual tuning properties, and molecular identities^1–6^. Despite this progress, how retinal output channels mature during development and how visual experience shapes this process remain incompletely understood.

In zebrafish, RGC axons project contralaterally to a defined set of retinorecipient brain areas known as arborization fields (AFs). Among these targets, the majority of RGC axons terminate in the largest AF10 or the optic tectum, the principal visual processing center^4,7–10^. The tectal neuropil is organized into nine layers, and individual RGCs choose a single tectal lamina as their target^8,11^. These stereotyped projection patterns collectively form approximately twenty RGC axonal projection classes (PCs)^4^. Some PCs occupy single tectal layers exclusively, as in the case of middle tectal lamina (SFGS2-5)^4^. Functional imaging of all RGC axons in response to behaviorally relevant visual stimuli revealed ten functional types^5,12^, with each tectal lamina generally encoding a restricted set of visual features. For instance, PC2 axons targeting superficial SO layer respond preferentially to small moving objects associated with prey capture^13^; PC4 axons terminating in SFGS1 layer respond selectively to motion direction underlying optokinetic and optomotor behaviors^14–16^; and PC8-11 axons terminating in deeper SFGS5/6 layers are associated with looming stimuli that evoke escape behavior^17^. Together, these observations suggest that each morphologically defined PC may generally correspond to a single functional type, contributing to dedicated visually guided behaviors^12^. However, this correspondence has largely been inferred from population-level analyses using pan-RGC labeling. Whether multiple functional response types coexist within the same tectal lamina, and whether single RGC transmit uniform tuning across all terminal branches remain open questions.

These issues are particularly important in the context of development. Zebrafish larvae begin exhibiting visually guided behaviors such as prey capture, optomotor responses, and looming-evoked escape by 5-7 days post-fertilization (dpf)^18–20^. Several of these behaviors continue to improve in efficiency and selectivity during the second week of development^21^, indicating ongoing maturation of visual circuit. At the same time, some RGCs undergo developmental changes in morphology and functional properties^22^, suggesting that retinal feature encoding is not fully fixed at the onset of behavior. However, it remains unresolved whether such changes reflect tuning refinement within individual neurons or population-level reorganization, as direct longitudinal tracking of single RGC across development has been lacking.

Normal visual development depends on both intrinsic molecular programs and environmental factors such as sensory experience. Neural plasticity driven by visual input has been extensively studied in cortical circuits in mammalian models^23,24^. In contrast, relatively little attention has been paid to how these principles operate within the retina, which has traditionally been regarded as a rigid, feedforward structure and a stable relay for downstream plasticity^25^. While early studies in the zebrafish visual system suggested that visual experience is dispensable for early structural and functional development^26–28^, more recent work has revealed critical roles of experience in shaping visual function and behavior^21,29–31^. Nonetheless, detailed analyses of functional plasticity at the level of specific retinal cell types remain limited^30^. Because RGCs form the final output of retinal processing, understanding whether distinct RGC subtypes undergo intrinsic or experience-dependent tuning changes is key to linking retinal development with adaptive visual behavior.

To address these gaps, genetically defined RGC populations offer a crucial entry point. In this study, we focus on RGCs expressing the transcription factor Satb2, a highly conserved regulator of chromatin organization and neuronal differentiation^32–35^. Previous studies have shown that Satb2 marks a restricted subset of RGC types, specifically direction-selective population, across multiple species^36–38^. In zebrafish, Satb2 is also expressed in ganglion cell layer and has been identified in three transcriptomic RGC types^6,34^. However, the morphological and functional properties of Satb2-positive RGCs in zebrafish remain uncharacterized.

Here, by combining *in vivo* two-photon calcium imaging with longitudinal single-cell tracking, we examined how tuning selectivity, spatial organization, and functional identity of RGC subtypes change from early to late larval stages. This approach reveals how diverse functional properties are represented onto individual layers of the optic tectum, and how intrinsic genetic programs and sensory experience interact to refine retinal output channels during development of behaviorally relevant visual circuits.

## Results

### Satb2-expressing RGC axons innervate three distinct layers of the optic tectum

To genetically label RGC subtypes marked by *satb2* gene, we generated transgenic lines in which the Gal4 transcription factor was inserted upstream of the *satb2* locus using CRISPR/Cas9-mediated knock-in. In the resulting *Tg(satb2:Gal4)* line, Gal4 expression was detected almost exclusively in RGCs within the brain, whose axons projected to the optic tectum (**Fig. 1a**). Additionally, we confirmed labeling around mouth as well as the retina, recapitulating the endogenous satb2 expression in pharyngeal arches and eye as previously observed at 48 hours post-fertilization (hpf)^30,39^. To verify that the transgenic line accurately recapitulates endogenous *satb2* expression, we compared GFP expression in *Tg(satb2:Gal4);Tg(UAS:GFP)* larvae with *satb2* and *Gal4* mRNA expression patterns detected by HCR in situ hybridization (**Fig. 1b**). The majority of RGCs labelled by the satb2:Gal4 driver also showed strong *satb2* mRNA signal. Conversely, *satb2* mRNA was largely restricted to Gal4-positive RGCs, suggesting that the transgenic line faithfully reflects the endogenous spatial pattern of *satb2* expression in the retina.

**Figure 1.**
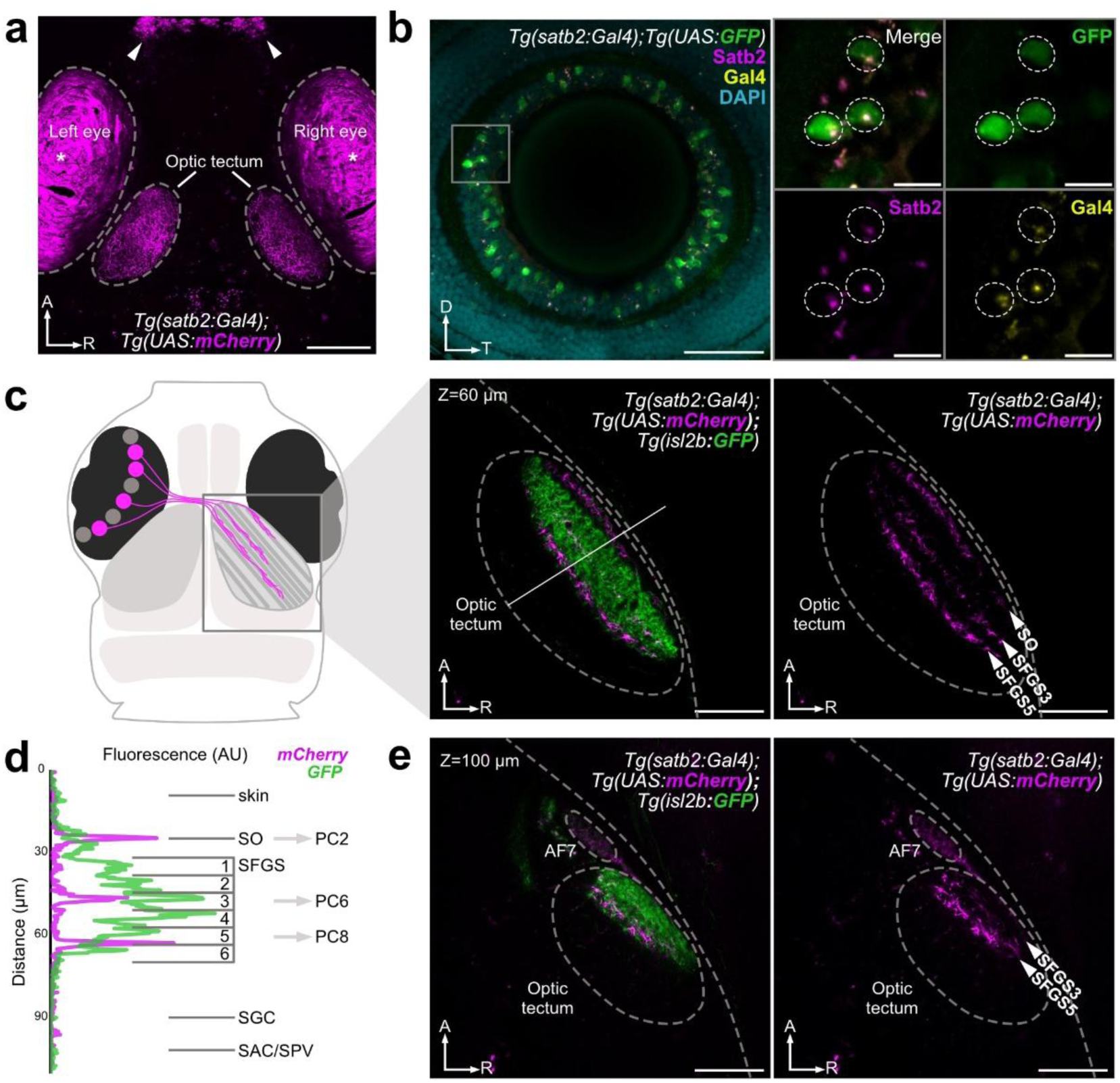
Axonal projection and retinal expression patterns of *satb2*- positive RGCs. **a)** Confocal Z-projection of a 7 dpf *Tg(satb2:Gal4);Tg(UAS:mCherry)* larva showing Satb2-positive RGC axons (magenta) in both optic tecta. White arrowheads mark expression around the mouth, and asterisks indicate eye autofluorescence. A, anterior; R, right. Scale bar, 100 µm. **b)** Composite confocal image of a 7 dpf *Tg(satb2:Gal4);Tg(UAS:GFP)* retina following HCR in situ hybridization for *satb2* (magenta) and *gal4* (yellow) mRNAs. Satb2-positive RGCs express GFP (green). Inset shows colocalization of *satb2* and *gal4* mRNA puncta within individual RGCs (dotted circles). Scale bars, 100 µm (left) and 10 µm (inset). **c)** Schematic (left) and confocal image (right) of *Tg(satb2:Gal4);Tg(UAS:mCherry);Tg(isl2b:GFP)* larva with grey box highlighting the right optic tectum shown in (c, e) at different depths. Isl2b-driven GFP labels all RGC axons (green), while Satb2-positive RGCs express mCherry (magenta). Right, the same image showing only Satb2-positive axons, which innervate three tectal laminae. Z = 60 µm refers to depths across the tectal Z-axis, with Z = 0 µm as the dorsal surface. Scale bar, 50 µm. **d)** Fluorescence intensity profile along the white line in (c) showing pan-RGC (GFP, green) and Satb2-positive (mCherry, magenta) axons. Three discrete magenta peaks mark Satb2-positive inputs to SO, SFGS3, and SFGS5 layers, corresponding to previously reported PC2, PC6, and PC8, respectively. **e)** Confocal image of the optic tectum at the deeper plane showing *satb2*-positive RGCs targeting AF7 at Z = 100 µm. Scale bar, 50 µm.

To determine the axonal projection pattern of Satb2-positive RGCs in the multilayered optic tectum, we performed confocal imaging of *Tg(satb2:Gal4);Tg(UAS:mCherry)* larvae crossed with *Tg(isl2b:GFP)*, in which all RGCs are labeled by GFP^40^ (**Fig. 1c**). Compared with the GFP-positive pan-RGC population, Satb2-positive axons target three distinct layers of optic tectum as well as AF7 (**Fig. 1d**, **e**). The fluorescence intensity analysis in the neuropil layers of the optic tectum revealed that the three layers correspond to SO, SFGS3 and SFGS5 layers (**Fig. 1d**). Comparing these information to previously established twenty PCs in zebrafish^4^ suggests that Satb2-positive axons terminating onto SFGS3 and SFGS5 layers correspond to PC6 and PC8, respectively. Both layers are innervated by dedicated inputs, e.g., having no collateral branches in other AFs^4^. Satb2-positive axons are found in SO and AF7 are likely to correspond to PC2, which terminate in SO layer while sending their collaterals to AF7 (**Fig. 1d**). Taking together, we conclude that Satb2-positive RGCs can be classified into at least three PCs, namely PC2, PC6, and PC8.

### Satb2-positive RGC axons are categorized into seven distinct functional clusters based on the visual response properties

To determine what kind of visual stimulus Satb2-positive RGCs respond to, we presented a battery of simplified visual features, each of which is relevant to a specific zebrafish behavior at 5-7 dpf. By 7 dpf, the density of synaptic contacts in the zebrafish retinotectal pathway has essentially reached its maximum^41,42^. At this stage, larvae are already capable of basic visually guided behaviors^43^. RGC responses to a monocular visual presentation were simultaneously recorded in the contralateral optic tectum via two-photon calcium imaging using 6-7 dpf larvae expressing cytosolic calcium indicator GCaMP6s in Satb2-positive RGCs (**Fig. 2a, b**). The stimulus set adopted and modified from previous study included visual stimuli that are relevant for behaviors of larval zebrafish: global luminance changes (dark and bright ramps and flashes) to probe for phototaxis; a small moving dot of 5°, which mimics the size of prey at the beginning of hunting behavior; a large moving dot of 30°, which approximates the size of prey immediately before the capture strike; fast and slow loom presented as an expanding disc at two different velocities to simulate an object approach that could trigger escape responses; moving gratings for detecting orientation- and direction-selective tuning^5^ (**Fig. 2c**). To identify RGC axons that responded to different stimuli in an unbiased manner, calcium time series were analyzed using a pixelwise analysis. We designed 18 different regressors for each of different stimulus variants and calculated a correlation score value (t-score) with these regressors for each pixel.

**Figure 2.**
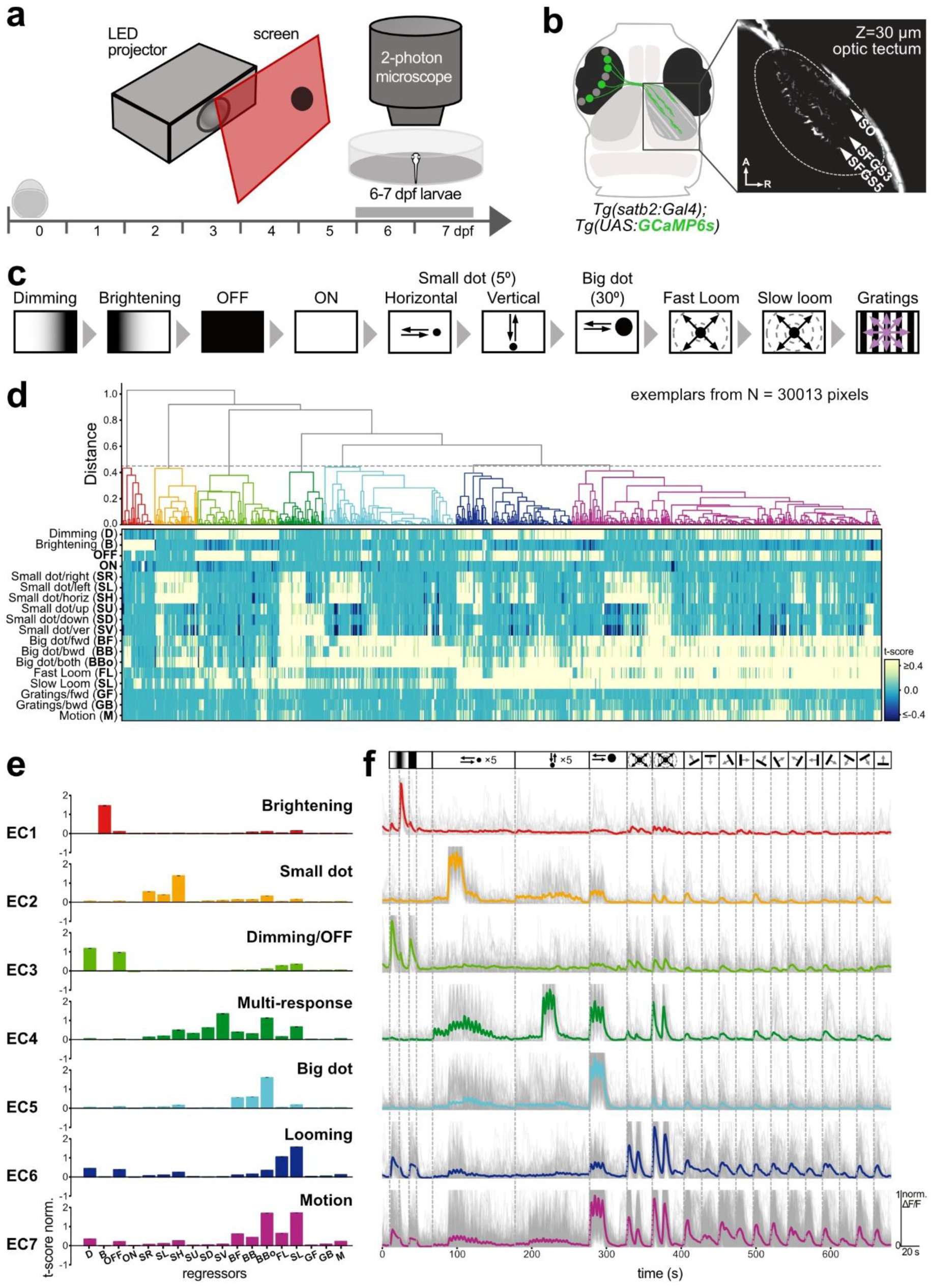
Functional imaging analysis of *satb2*-positive RGCs at early larval stage. **a)** Experimental set up for functional imaging. *In vivo* calcium imaging was performed using a two-photon microscope on 6-7 dpf *Tg(satb2:Gal4);Tg(UAS:GCaMP6s)* transgenic zebrafish larvae. Visual stimulus was projected on white diffusive screen using the red channel of LED projector. Contralateral optic tectum was simultaneously imaged as visual stimuli were presented monocularly. Schematic timeline indicates the developmental stages, marking the period of functional data acquisition at 6-7 dpf (grey bar). **b)** Two-photon calcium imaging of 7 dpf *Tg(satb2:Gal4)* larvae expressing cytosolic GCaMP6s in RGCs found in SO, SFGS3, and SFGS5 layers. The optic tectum was imaged at several depths across Z-plane and Z = 0 is defined as beginning of optic tectum from dorsal most side. **c)** Visual stimulation protocol consisted of dimming and brightening; dark and bright flash; a small horizontally moving dot (5° presented at five equally spaced elevations across the vertical axis of the screen); a small vertically moving dot (5° presented at five equally spaced azimuths across the horizontal axis of the screen); a big moving dot presented horizontally (30° in forward and backward directions with two repetitions); looming stimuli presented in two velocities (a fast and a slow loom); grating stimulus consisting of black gratings moving in 12 equally spaced angular directions. **d)** Hierarchical clustering dendrogram and corresponding heatmap of response profiles from N = 30,013 pixels imaged from Satb2-positive RGC axons in the optic tectum (N = 6 fish). A grey dotted line indicates a chosen distance threshold of 0.45, which yielded seven functional clusters named as Early Clusters (ECs) with a distinct color code. The heatmap shows responses (t-score) of each pixel to 18 regressor types listed and abbreviated along y axis. **e)** Bar plots show the relative contribution of each stimulus feature (x-axis: 18 regressors; y-axis: normalized t-score) to the clusters’ responses. EC1 is associated with brightening/red color; EC2 with small dot/yellow; EC3 with dimming and OFF/green; EC4 shows a multi-response pattern/dark green; EC5 with big dot/blue; EC6 with fast and slow loom/purple; and EC7 shows mixed responses to dimming, OFF, big dot, fast loom, slow loom, and gratings/magenta. For abbreviation of the 18 regressors, see (d). **f)** Normalized activity traces (ΔF/F0) for each early cluster (EC1-7). All corresponding exemplars that fall into the cluster are plotted in grey, while the average of all pixels within a cluster are plotted with bold colored lines, according to the color code defined in (d). Stimulus epochs are indicated above the traces and dotted vertical lines represent the onset of the corresponding stimulus.

To better classify functional response types, we performed hierarchical clustering of representative response patterns using affinity propagation. The fish analyzed at the developmental stage of 6-7 days is referred as 7 dpf. A pixel-wise regressor and cluster analysis resulted in the dendrogram and corresponding heatmap for 704 exemplars with the total of 30,013 responsive pixels from six fish at 7 dpf (**Fig. 2d**). Classification of all fish yielded seven major clusters (also referred as Early Cluster or EC) with a distinct set of responses to the presented battery of visual stimuli (**Fig. 2e**, **f**; see also **Methods**). Certain clusters exhibited strong selectivity for one or two specific stimuli, such as brightening, small dots, dimming/OFF, or big dots, corresponding to EC1, EC2, EC3, and EC5, respectively. In contrast, the remaining clusters comprised pixels responsive to a broad range of multiple visual stimuli. For instance, EC4 responded to small dots, big dots, and slow/fast looming stimuli and was annotated as multi-response; EC6 responded to dimming/OFF and slow/fast looming stimuli and was annotated as looming; and EC7 responded to small dots, big dots, slow/fast looming stimuli, and moving gratings and was annotated as motion response. (**Fig. 2e**, **f**). Among these clusters, EC5, EC6, EC7 and EC3 constituted around three quarters (81.9%), while EC1, EC2 and EC4 each accounted for less than 10% of the total population (**Extended Data Fig. 1a**). Similar distribution pattern was consistently observed across six different fish at 7 dpf (**Extended Data Fig. 1b**).

To assess whether the functional response profiles are internally consistent within each cluster and also distinctive across different clusters, we conducted a pairwise Pearson correlation analysis between all responsive pixels across the dataset (N = 30,013 pixels). This analysis was performed for pixel-level and cluster-average-level comparisons (**Extended Data Fig. 1c, d**). Overall, we found high correlation between pixels within the same cluster (**Extended Data Fig. 1c**), resulting in strong inter-cluster similarity with block structures along the diagonal for each cluster in the cluster-averaged correlation matrix (r = 1 for same cluster comparison, **Extended Data Fig. 1d**). When comparing inter-cluster correlations, we found that EC1 (brightening), EC2 (small dot), and EC3 (dimming/OFF) show generally negligible or negative correlations to other clusters, indicating their distinct tuning. By contrast, EC7 (motion), unlike EC1 and EC2, shares notable correlations with EC5 (big dot) and EC6 (looming), both around r = 0.7 at the cluster-average-level, suggesting partial response overlap. These analyses not only validate the robustness of clustering but also highlight both discrete and graded organization of early visual response types among Satb2-positive RGCs. Taken together, these findings indicate that Satb2-positive RGCs respond to a variety of behaviorally relevant visual stimuli.

### Different functional response types of Satb2-positive RGCs are topographically organized and coexist in single tectal layers

To examine the spatial distribution of Satb2-positive RGC functional responses, we mapped visually responsive pixels from seven functional clusters at 7 dpf to their corresponding positions in the optic tectum of the standard zebrafish brain using the ANTs registration workflow, as detailed in the Methods section (**Fig. 3a**, **b**; see also **Extended Data Fig. 2a**). Overlaying GCaMP6s signals from six fish confirmed that axon terminals of Satb2-positive RGCs were precisely aligned within three discrete tectal layers (**Extended Data Fig. 2b**).

**Figure 3.**
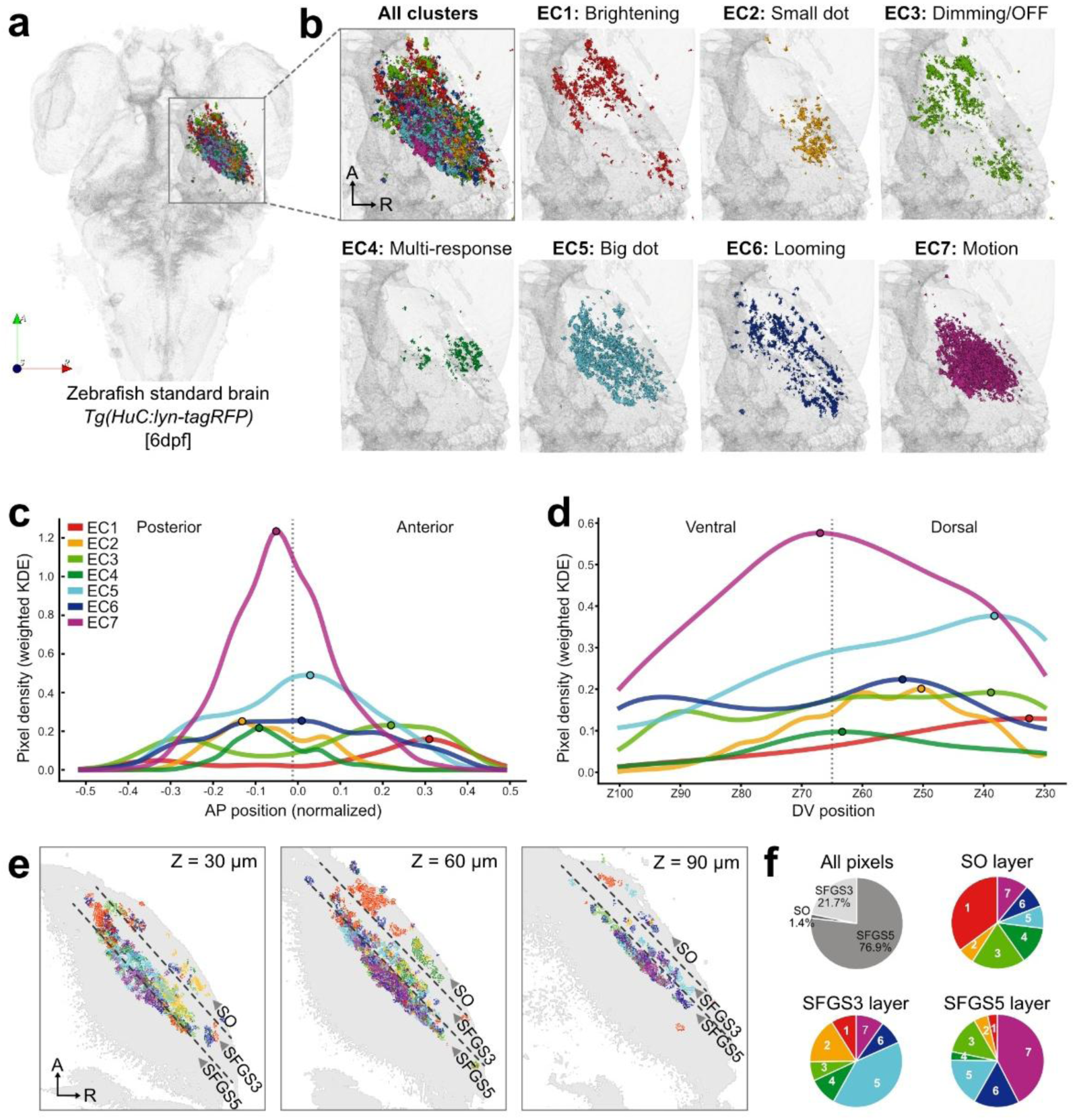
Anatomical characterization of Satb2-positive RGCs using clustering analysis. **a)** Top view of the 6 dpf zebrafish standard brain (*Tg(HuC:lyn-tagRFP)* from mapZebrain atlas) with an inset highlighting the region where functional Satb2-positive RGC terminals of all seven ECs from six fish at 7 dpf are registered. A, anterior; R, right; S, superficial. **b)** Upleft panel shows spatial distribution of all EC pixels registered to the standard brain. Remaining panels show individual clusters (EC1-7) displayed separately and color-coded as defined in Fig. 2e. A, anterior; R, right. **c)** Distribution of cluster pixels along the anterior-posterior (AP) axis at 7 dpf. Weighted kernel density estimates (KDEs) of normalized AP positions (-0.5 = posterior, 0.5 = anterior) are plotted for each cluster. Dots along the line indicate AP mode (position of maximal pixel density) for each cluster. **d)** Distribution of cluster pixels along the dorso-ventral (DV) axis at 7 dpf. Weighted KDEs of normalized DV positions (Z100 = ventral, Z30 = dorsal) are plotted for each cluster. Dots along the line indicate DV mode (position of maximal pixel density) for each cluster, with DV coordinates systematically assigned from 0.0 = Z100 (ventral) to 0.7 = Z30 (dorsal) in 0.1 increments per plane. **e)** Cross-sectional images (Z-slices) at three depths (Z = 30 µm, 60 µm, 90 µm) showing laminar distribution of registered LC pixels following color code complementing (b). A, anterior; R, right. **f)** Cluster composition within each tectal laminae at 7 dpf. Upleft: overall distribution of all EC pixels across three tectal laminae, showing preferential targeting of SFGS5. Remaining pie charts show proportions of ECs contributing to each of three layers.

Cluster-specific mapping revealed distinct spatial biases across the optic tectum (**Fig. 3b**). EC1 (brightening) and EC3 (dimming/OFF) terminals were located predominantly in the anterior tectum, whereas EC2 (small dot) and EC4 (multi-response) terminals were shifted toward posterior regions. In contrast, the clusters with the larger pixel counts including EC5 (big dot), EC6 (looming), and EC7 (motion) were broadly distributed across the anterior-posterior axis. To quantify these spatial trends, we analyzed pixel distributions along the anterior-posterior (AP) and dorso-ventral (DV) axes (**Fig. 3c, d**). Along the AP axis, we confirmed the anterior bias of EC1 and EC3 (AP mode or highest estimated pixel density along AP axis < 0.2), the posterior enrichment of EC2 and EC4 (AP mode around -0.1), and the broader distributions of EC5, EC6, and EC7, with EC7 distribution peaking in the middle part of optic tectum. Along the DV axis, EC4 and EC7 were concentrated in the middle position of the imaged volume (DV mode or highest estimated pixel density along DV axis <0.4), whereas other clusters were distributed from middle to dorsal part of tectal neuropil (DV mode > 0.4). These topographic patterns indicate a non-uniform representation of visual feature selectivity along the retinotopic axes, consistent with the asymmetric distribution of RGC subtypes in the retina^1,5,44^.

We next closely examined how different ECs are distributed within individual tectal layer. Since each of SO, SFGS3, and SFGS5 layers is innervated by a single unique projection class^4^, we asked whether a single layer represents one functional response or accommodates multiple response types. Results from single representative planes show that none of the layers were functionally homogeneous (**Fig. 3e**; **Extended Data Fig. 2c**). In fact, each of three tectal layers contained responses from all ECs represented in varying proportions, and this heterogeneity persisted along the dorso-ventral planes (**Fig. 3e**, **f**). Across all layers, functionally active Satb2-positive RGC axon terminals were most abundant in SFGS5, which contained nearly three-quarters of all responses, with the remainder distributed between the SFGS3 and SO layers (**Fig. 3f**).

Taken together, these findings indicate that Satb2-positive RGCs respond to a variety of behaviorally relevant visual stimuli, not only across different layers of the optic tectum but also within individual layers. This suggests that multiple functional types coexist within a layer that is innervated by a dedicated input from a single PC (i.e., SFGS3 and SFGS5).

### Multiple response types observed within a single tectal layer originate from RGCs, each of which exhibit a single, distinct response pattern

The presence of multiple response clusters within a single tectal lamina raises at least two possibilities (**Fig. 4a**). Hypothesis 1 proposes that individual RGCs possess uniform responses across all axon terminals, and that functional diversity within a lamina arises from the convergence of differentially tuned RGC types. Hypothesis 2 predicts that a single RGC may exhibit multiple functional responses across its terminal branches, generating intralaminar heterogeneity, similar to reports of multiple branch-specific tunings in bipolar cell axons^45^. Distinguishing between these possibilities requires resolving response identity at the level of individual RGCs rather than at the population average.

**Figure 4.**
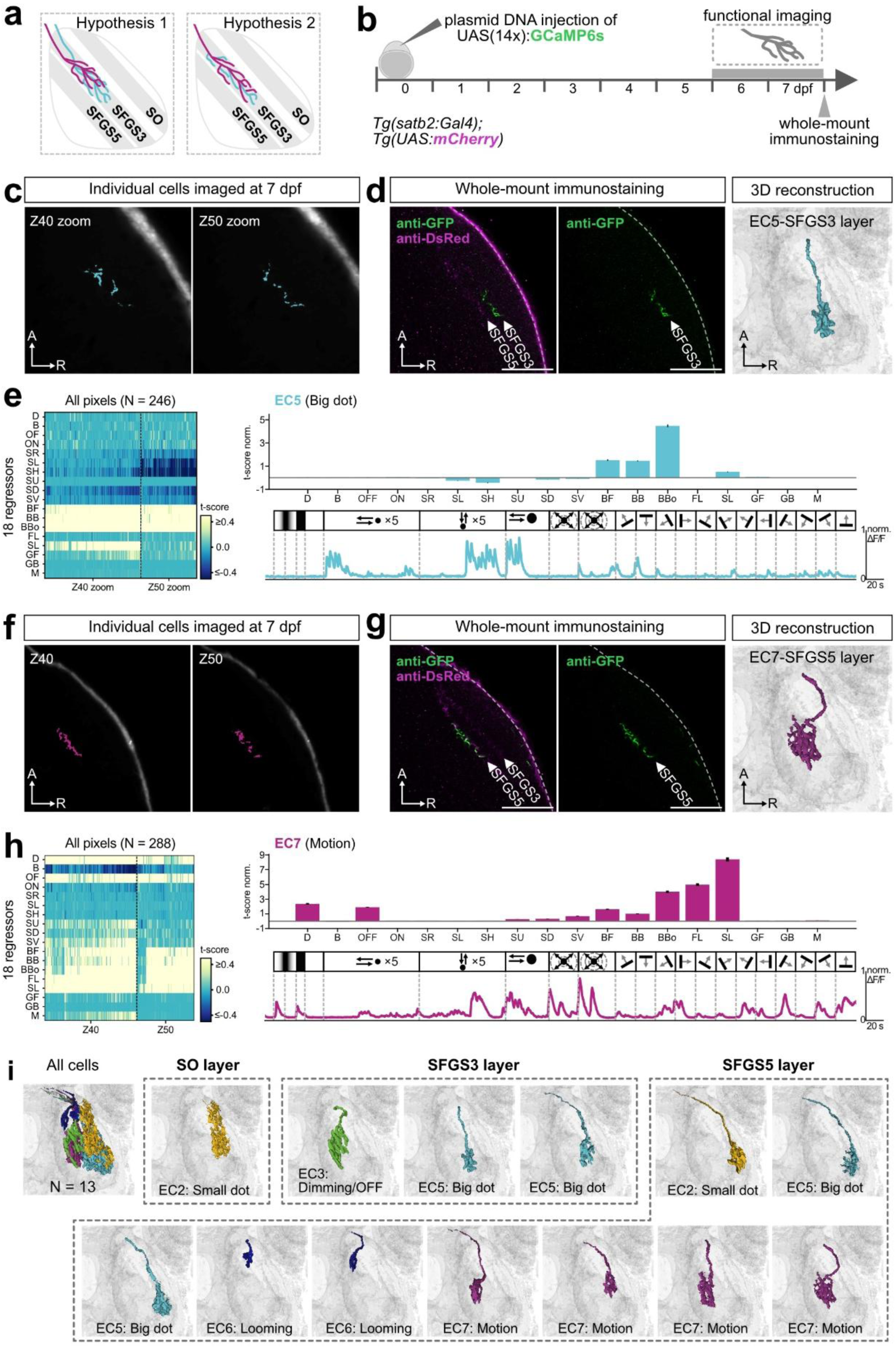
Sparse labeling of Satb2-positive RGCs to show spatial organization of functionally distinct RGC types. **a)** Two hypothetical models for the presence of multiple functional response types within the same retinorecipient tectal layer. *Hypothesis 1:* Multiple RGCs, each exhibiting a homogeneous response across all axonal branches, converge onto the same tectal layer. *Hypothesis 2:* A single RGC possesses distinct axonal branches with heterogeneous response profiles that terminate in a single tectal layer. **b)** Experimental workflow. Sparse labeling of Satb2-positive RGCs was achieved by injecting UAS(14x):GCaMP6s plasmid DNA into *Tg(satb2:Gal4);Tg(UAS:mCherry)* embryos at singe-cell stage. Functional calcium imaging was performed at 6-7 dpf, followed by whole-mount immunohistochemistry to match functional responses to terminal anatomy. **c)** Two-photon images of sparsely labeled terminals from individual RGCs at 7 dpf across two Z-planes (Z40, Z50). A, anterior; R, right. **d)** Post-hoc whole-mount immunostaining of the imaged cell in (c). GFP labels the injected GCaMP6s-expressing RGC; DsRed labels the three Satb2-positive recipient layers (SO, SFGS3, SFGS5). 3D reconstruction confirms that this neuron arborizes selectively within the SFGS3 lamina. Color indicates cluster assignment derived from (e). Scale bar: 50 µm. **e)** Functional response properties of the RGC shown in (c-d). Left: pixel-wise response heatmap (t-score) across 18 regressors from two imaging planes (Z40, Z50). Right: average response profiles shown as bar plot and normalized activity traces (the average of all pixels) (ΔF/F0), indicating strong preference for bog-dot stimuli, classified as EC5 (big dot). **f-g)** Same layout as (c-d) for another sparsely labeled RGC terminating within the SFGS5 layer. 3D reconstruction confirms laminar specificity. Color denotes functional cluster from (h). A, anterior; R, right. Scale bar, 50 µm. **h)** Functional response map of cell shown in (f-g). Left: pixel-wise t-score matrix for 18 regressors across two imaging planes. Right: average response profiles shown as bar plot and normalized activity traces (the average of all pixels) (ΔF/F0), showing strong activation to multiple motion-related stimuli, classified as EC7 (motion). **i)** Summary of reconstructed Satb2-positive RGC morphologies (N = 13) organized by laminar position. Each example represents one cell, color-coded by functional cluster identity. SO layer: EC2 (small dot). SFGS3 layer: EC3 (dimming/OFF), EC5 (big dot). SFGS5 layer: EC2 (small dot), EC5 (big dot), EC6 (looming), EC7 (motion).

To test these scenarios, we sparsely expressed GCaMP6s in Satb2-positive RGCs by injecting UAS(14x):GCaMP6s plasmid DNA into *Tg(satb2:Gal4);Tg(UAS:mCherry)*, thereby labeling only a few RGCs for functional analysis at 7 dpf (**Fig. 4b**). Two-photon recordings of a representative sparsely labeled RGC showed that all terminal branches, imaged across several Z-planes, exhibited highly homogeneous calcium responses (**Fig. 4c**). Post-hoc whole-mount immunostaining confirmed that all arbors were confined to the SFGS3 layer, and 3D reconstruction further verified that these terminals belonged to a single RGC with multiple branches converging into a long axon (**Fig. 4d**). A more detailed pixel-wise analysis revealed that response preference of all pixels across planes for moving big-dot stimuli was consistent with EC5 tuning derived from population-level data at 7 dpf, as reflected in heatmap, average bar plot and activity trace (**Fig. 4e**). Another example of homogeneous response preference is shown for a single RGC whose axon terminals reside in the SFGS5 layer and are classified into the EC7 (motion-sensitive) cluster (**Fig. 4f-h**).

We extended this approach to a total of 13 sparsely labeled RGCs, reconstructing their axonal projection in the optic tectum and assigning response identity to each cell (**Fig. 4i**). In every case, terminals belonging to the same RGC exhibited a single, internally consistent functional profile, with no evidence of branch-specific tuning within a single neuron. Consistent with the relative distribution of each tectal lamina, we predominantly targeted SFGS5-projecting neurons which belonged to EC2 (small dot), EC5 (big dot), EC6 (looming), and EC7 (motion). In contrast, only one RGC was sparsely labeled in SO layer with an EC2 response, and a few individual RGCs were identified in SFGS3 layer with EC3 (Dimming/OFF) or EC5 response patterns.

Together, these data support Hypothesis 1, demonstrating that functional diversity within a given tectal layer arises from the convergence of multiple RGC types rather than from mixed selectivity emerging within individual axons. Single RGCs therefore operate as functionally coherent units at their axon terminals, each conveying a dominant set of visual features, while lamina accumulates diversity through the parallel arrival of distinct response-specific RGC types.

### Developmental maturation of functional response properties of Satb2-positive RGCs

By the second week of development around 10 dpf, zebrafish larvae display more sophisticated visually guided behaviors, such as efficient prey capture^21^. These behavioral refinements coincide with continued retinal maturation, including elaboration of RGC dendritic stratification and synaptic connectivity by interneurons such as bipolar and amacrine cells^46,47^. Previous work has shown that functional properties of some RGC subtypes, such as orientation-selective cells, also undergo developmental refinement during this period^22^. To determine whether the response types observed in Satb2-positive RGCs at 7 dpf are stably maintained or developmentally reorganized, we performed functional imaging at 10-11 dpf (hereafter referred to as 11 dpf; **Fig. 5a**).

**Figure 5.**
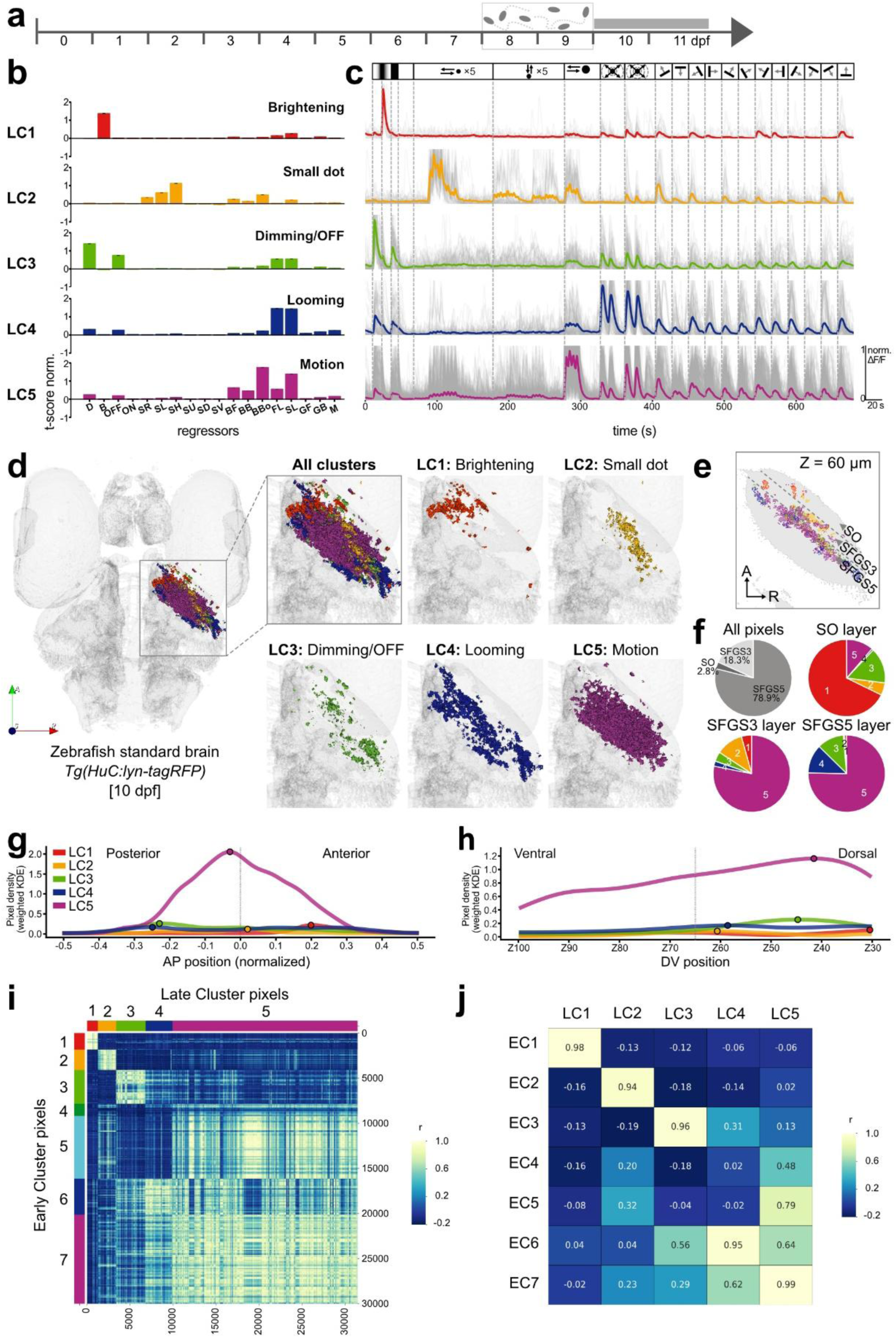
Functional and anatomical characterization of Satb2-positive RGCs over development. **a)** Schematic timeline indicates the developmental stages, marking the period of functional data acquisition at 10-11 dpf (grey bar). Larvae were fed with paramecia starting from 7 dpf and until functional imaging (moving paramecia in grey). N = 6 fish. **b)** Bar graphs showing the average normalized response weights of five functional RGC Late Clusters (LC1-5) at 10-11 dpf to 18 visual stimulus regressors. LC1 is primarily responsive to brightening, LC2 to small dot, LC3 to dimming/OFF, LC4 to looming stimuli, and LC5 to motion stimuli with color-code inherited from early clusters with similar functional response types. **c)** Normalized calcium activity traces (ΔF/F₀) of LC1-5. Each colored trace is an average of the cluster activity, with individual pixel exemplars shown in grey. **d)** Spatial distribution of functionally defined LC pixels registered to the 10 dpf standard brain of *Tg(HuC:lyn-tagRFP)*. The inset and enlarged panel highlight the region where functional Satb2-positive RGC terminals of all five LCs from six fish at 11 dpf are registered. Remaining panels show individual clusters (LC1-5) displayed separately and color-coded as defined in (b). A, anterior; R, right; S, superficial. **e)** Cross-sectional Z-slice at the depth of Z = 60 µm, showing laminar distribution of registered LC pixels. A, anterior; R, right. **f)** Functional pixel composition of all LC pixels across all tectal laminae and cluster-classified pixels within each tectal laminae at 11 dpf. **g)** Distribution of cluster pixels along the anterior-posterior (AP) axis at 11 dpf. Weighted kernel density estimates (KDEs) of normalized AP positions (-0.5 = posterior, 0.5 = anterior) are plotted for each cluster, color-coded as in (b). Dots along the line indicate AP mode for each cluster. **h)** Distribution of cluster pixels along the dorso-ventral (DV) axis at 11 dpf. Weighted KDEs of normalized DV positions (Z100 = ventral, Z30 = dorsal) are plotted for each cluster. Dots along the line indicate DV mode for each cluster, with DV coordinates systematically assigned from 0.0 = Z100 (ventral) to 0.7 = Z30 (dorsal) in 0.1 increments per plane. **i)** Pixel-to-pixel correlation heatmap showing hierarchical similarity between all individual pixels, ordered first by early cluster identity (EC1-7, vertical) and by late cluster identity (LC1-5, horizontal). Color coding along the axes represents early (left) and late (top) cluster assignments. Each matrix entry corresponds to the Pearson correlation (r) of stimulus response profiles between two pixels. Diagonal blocks indicate high within-cluster similarity, while off-diagonal patterns reflect transitions or overlaps between early and late functional clusters. **j)** Cluster-to-cluster similarity matrix based on Pearson correlation coefficients (r) showing pairwise similarity between each cluster of ECs and LCs.

*De novo* hierarchical clustering of responses at 11 dpf identified five distinct groups, termed Late Clusters (LC1-5), using the same analytical parameters applied at 7 dpf (**Fig. 5b**, **c**; **Extended Data Fig. 3a**). Comparison of mean response profiles revealed that LC1-3 closely resembled EC1-3, indicating stable preservation of brightening-, small dot-, dimming/OFF-responsive types across development. LC4 and LC5 shared hallmark features of early looming (EC6) and motion (EC7) responses, respectively, but displayed broader tuning compared to their early counterparts. For example, LC4 responded robustly to both slow and fast looming stimuli, whereas its early counterpart EC6 showed a stronger preference for slow loom (**Fig. 2g**, **5c**). Furthermore, motion responses to moving gratings were more prominent in LC4 than in EC6, suggesting an emergence of more complex tuning properties at later stages. In contrast, EC4 (multi-response) and EC5 (big dot) did not persist as distinct clusters at 11 dpf, indicating that these early types may be pruned or subsumed into later functional groups.

While the retinotopic and laminar organization of Satb2-positive RGC responses at 11 dpf closely resembled that at 7 dpf, the composition of functional clusters within each layer underwent marked redistribution. Anatomical mapping revealed that spatial locations of late clusters largely recapitulated early-stage patterns (**Fig. 5d**): LC1 (brightening) remained confined to the anterior optic tectum, while LC3 (dimming/OFF) and LC4 (looming) localized to more posterior positions. In contrast, LC2 (small dot) and LC5 (motion) spanned the full anterior-posterior axis, with LC5 occupying a substantially larger area. Multiple clusters were still observed within the same tectal lamina (**Fig. 5e**), indicating that each anatomical layer continued to host diverse functional response types even at the later developmental stages.

A striking shift emerged in relative abundance of response types (**Extended Data Fig. 3b**, **c**). LC5 accounted for 68.5% of all classified pixels at 11 dpf, more than double the fraction of its putative early-stage precursor, EC7 (32.9 %, **Extended Data Fig. 1a**, **3b**). This dominance was evident across layers, with LC5 representing around 75% pixels in both SFGS3 and SFGS5 (**Fig. 5f**). In contrast, each of LC1-4 remained similar in overall proportions to their early counterparts, although LC1 with selective brightening response was especially enriched in superficial SO layer at the late stage. Quantitative retinotopic analysis demonstrated that the overall AP and DV positions of clusters were preserved, confirming stable laminar targeting and spatial organization across development. However, unlike other clusters, LC5 showed a pronounced expansion. Despite occupying a similar central-anterior region along the AP axis as EC7 (**Fig. 5g**), LC5 shifted from ventral side along the DV axis as EC7 at 7dpf toward the dorsal side by 11 dpf (**Fig. 5h**). Overall, correlation analysis further indicated stronger separation among clusters at 11 dpf, with reduced pairwise correlations compared to 7 dpf (**Extended Data Fig. 3d**, **e**), suggesting increased functional specialization as circuits mature.

### Developmental comparison of functional cluster identity and distribution between 7 and 11 dpf

To directly relate early functional response types to their later counterparts, we compared the 7 dpf early clusters (ECs) and the 11 dpf late clusters (LCs). Pearson correlation analysis was performed to both at the pixel level (**Fig. 5i**) and at the cluster level based on their mean response profiles (**Fig. 5j**). These analyses revealed a strong one-to-one correspondence between EC1-3 and LC1-3 (cluster-wise Pearson r > 0.9), respectively, indicating robust preservation of brightening, small dot, and dimming/OFF response types across development. EC6, which responded predominantly to looming stimuli, showed high similarity to LC4 (r = 0.95), strongly supporting its role as the developmental precursor of the late looming-sensitive group. Likewise, EC7 aligned closely with LC5 (r = 0.99), demonstrating that motion sensitivity is stably maintained from early to late stages.

On the other hand, EC4 (multi-response) and EC5 (big-dot) did not display a clear one-to-one correspondence with any of the late clusters. EC5 exhibited a moderate correlation with LC5 (r = 0.79), suggesting that big-dot-responsive cells may be potentially integrated into broader motion-related response characteristic of LC5. These trends were consistent across both pixel- and cluster-level correlation analyses.

We then asked how these population-level functional transformations are brought about at the cellular level. To test whether individual cells undergo corresponding changes in response profiles, we performing longitudinal functional imaging of sparsely labeled Satb2-positive RGC axons first at 7 dpf and again at 11 dpf. Whole-mount immunostaining was subsequently used to validate the anatomical identity of each cell (**Fig. 6a**). This approach allowed direct tracking of response dynamics within the same neuron across developmental stages.

**Figure 6.**
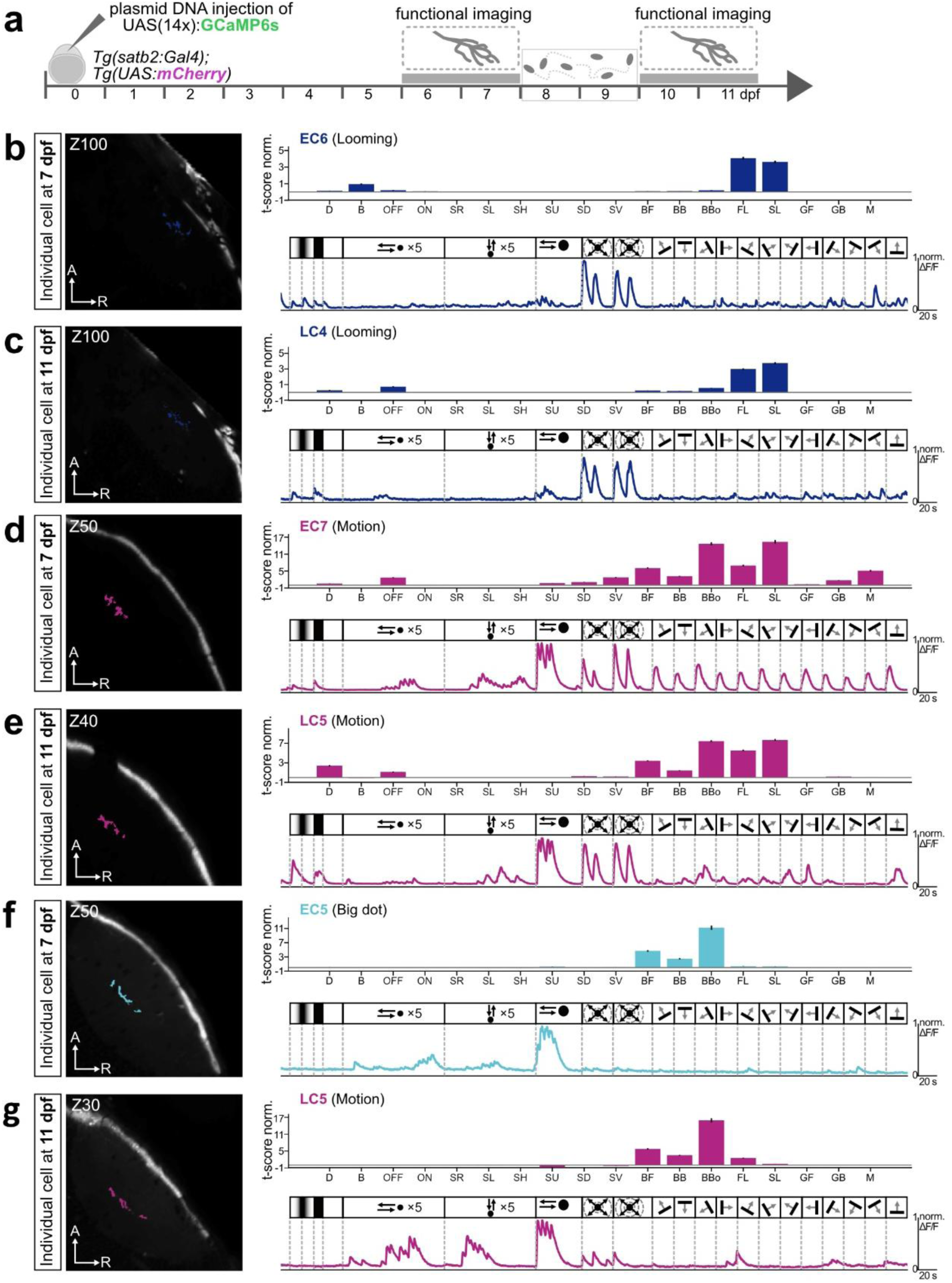
Longitudinal tracking of Satb2-positive RGCs reveals stable and dynamic changes in functional responses between early and late development. **a)** Experimental workflow for repeated two-photon imaging of sparsely labeled Satb2-positive RGCs. *Tg(satb2:Gal4);Tg(UAS:mCherry)* embryos were injected with UAS(14x):GCaMP6s plasmid DNA, enabling re-identification of single RGC terminals and comparison of functional response properties at 7 and again 11 dpf. In all panels, Z-depth indicates imaging plane (µm), RGC terminals are color-coded by cluster identity, and ΔF/F₀ traces correspond to the stimulus sequence shown above. A, anterior; R, right **b-c)** Example of an RGC that maintains a looming-selective response over development. At 7 dpf (b), the cell is classified as EC6: Looming, and at 11 dpf (c) it remains looming responsive (LC4: Looming), indicating a stable functional identity (N = 3). **d-e**) A motion-responsive RGC retains its identity through development. At 7 dpf (d) the cell corresponds to EC7: Motion, and at 11 dpf (e) it maps to LC5: Motion, reflecting persistence of motion tuning (N = 6). **f-g)** Example of a response transition between developmental stages. An RGC initially categorized as EC5: Big dot at 7 dpf (f) shifts to a motion-dominated classification (LC5: Motion) at 11 dpf (g), demonstrating functional reorganization in late development (N = 2).

At the single-cell level, we observed clear continuity in functional identity. EC6 (looming) cells at 7 dpf reliably developed into LC4 (looming) at 11 dpf (**Fig. 6b**, **c**; N = 2 cells). Similarly, EC7 (motion) cell matured into LC 5 (motion) (**Fig. 6d**, **e**; N = 7 cells). Closer observation revealed that a cell broadly tuned to all directions of grating movement at 7 dpf became selectively responsive to backward-moving gratings at the later stage (**Fig. 6e**). This is consistent with the emergence of direction-selective, backward-tuned responses during later stages of development (**Extended Data Fig. 4a**, **c**), while orientation-selective responses remained unchanged (**Extended Data Fig. 4b**, **d**). It is worth noting that orientation-selective RGCs have been reported to undergo functional refinement in zebrafish^22^, and such changes may occur in other RGC subtypes not captured by Satb2 labeling. Thus, our data suggest that Satb2-positive RGCs specifically contribute to the emergence of direction-selective motion processing, a property likely important for the maturation of visually guided behaviors such as predator avoidance and/or prey capture.

Notably, single RGCs classified as EC5 (big dot) at 7 dpf, though not present as a distinct cluster at 11 dpf, were reassigned as LC5 (motion) at the late stage (**Fig. 6f, g**; N = 4 cells). Closer observation of the bar plots and activity traces shows that cell initially exhibiting a selective response to the big-dot stimulus at 7 dpf begin to develop clear responses to looming stimuli, particularly fast looming, by 11 dpf. These transitions provide a cellular explanation for the expansion of LC5 at the population level and suggest that EC5 neurons can be repurposed into the broader motion-responsive pool during late development.

Together, these findings suggest that while several RGC response types remain stable across development, others undergo functional refinement and integration into later clusters. In particular, the maturation and expansion of motion-selective RGCs could contribute to the emergence of a more specialized and behaviorally relevant functional architecture by 11 dpf.

### Temporary visual deprivation modifies Satb2-positive RGC functional clustering

Larvae at 10-11 dpf were fed paramecia between 8-10 dpf since the yolk-derived nutrients are depleted by this stage and feeding becomes essential for survival. This introduces the possibility that the functional changes observed between 7 and 11 dpf may arise from: (1) intrinsic developmental maturation, (2) visual experience associated with prey detection or surrounding environment, or (3) the act of prey consumption itself. To assess the contribution of visual experience to RGC functional maturation, we designed an experiment where larvae were reared in darkness from 7 to 11 dpf, while still being fed with paramecia. Larvae were kept under aluminum foil to exclude light exposure (**Fig. 7a**). Under this condition, the larvae still consumed paramecia, as judged by their stomach while fed and unfed, probably using other senses than vision, such as lateral line or olfactory systems (**Fig. 7b**). Functional imaging and hierarchical clustering at 11 dpf in dark-reared fish yielded the same five functional clusters (dark-reared late cluster or DRLC) as those in normally reared fish (**Fig. 7c**, **d**; **Extended Data Fig. 5a**). Anatomical maps of each DRLC, their fractions and internal cluster correlation were also remarkably similar to LCs, suggesting a negligible impact of visual exposure on overall functional maturation and localization (**Fig. 7e**, **f**; **Extended Data Fig. 5b-e**).

**Figure 7.**
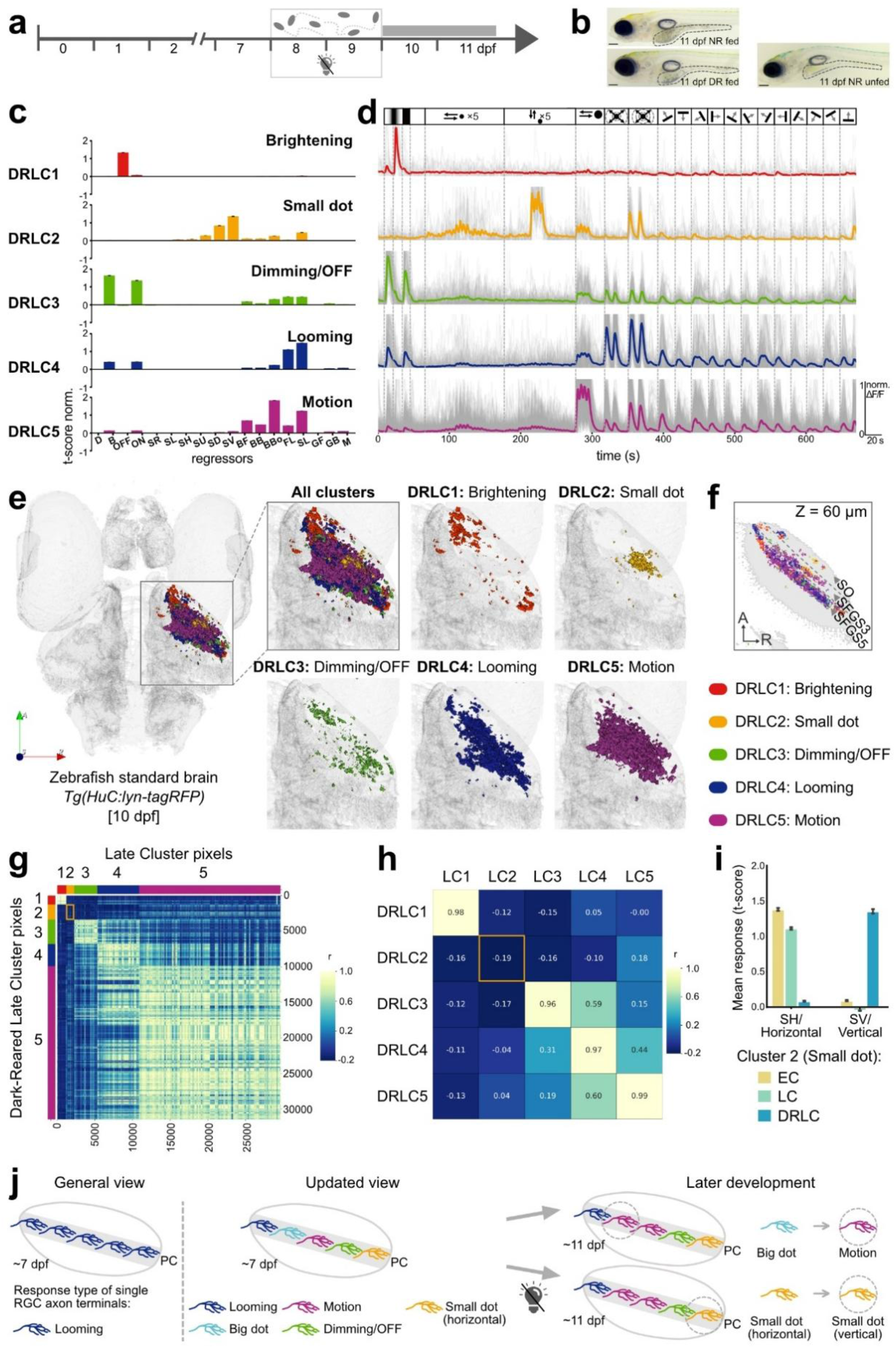
Functional and anatomical characterization of Satb2-positive RGCs over development and temporary visual deprivation. **a)** Schematic timeline indicates the developmental stages, marking the period of functional data acquisition at 10-11 dpf. Larvae were fed with paramecia and covered with aluminum foil for dark rearing starting from 7 dpf and until functional imaging. N = 6 fish. **b)** Images of fed (normal-reared 11 dpf), fed (dark-reared 11 dpf) and unfed (normal-reared 11 dpf) larval zebrafish with grey dotted line indicated stomach. Zebrafish fed with paramecia has accumulation appearing as grey in the stomach compared to unfed. **c)** Bar graphs showing the average normalized response weights of five functional RGC Dark-Reared Late Clusters (DRLC1-5) at 11 dpf to 18 visual stimulus regressors. DRLC1 is responsive to brightening, DRLC2 to small dot, DRLC3 to dimming/OFF, DRL4 to looming stimuli, and DRLC5 to motion stimuli. **d)** Normalized calcium activity traces (ΔF/F₀) of DRLC1-5. Each colored trace is the average of the cluster activity with individual pixel exemplars shown in grey. **e)** Spatial distribution of functionally defined DRLC pixels registered to the 10 dpf standard brain of *Tg(HuC:lyn-tagRFP)*. The inset and enlarged panel highlight the region where functional Satb2-positive RGC terminals of all five DRLCs from six fish at 11 dpf are registered. Remaining panels show individual clusters (DRLC1-5) displayed separately and color-coded as defined in (b). A, anterior; R, right; S, superficial. **f)** Cross-sectional Z-slice at the depth of Z = 60 µm, showing laminar distribution of registered DRLC pixels. A, anterior; R, right. **g)** Pixel-to-pixel correlation heatmap displaying Pearson correlation coefficient (r) of all individual pixels of LC with DRLC at 11 dpf. Each block along the diagonal reflects within-cluster coherence, while off-diagonal blocks indicate inter-cluster similarity. Pixels are ordered by cluster assignment. **h)** Cluster-to-cluster similarity matrix showing Pearson correlation coefficients (r) for mean response patterns between each cluster of LC and DRLC. **i)** Bar plots summarizing the mean regressor t-scores (± SEM) of two regressors: SH (small dot moving horizontally) and SV (small dot moving vertically). Responses are shown for Early Clusters (EC, yellow), Late Clusters (LC, teal), and Dark-Reared Late Clusters (DRLC, blue). **j)** Conceptual model of developmental functional organization and refinement of Satb2-positive RGC outputs. Left, general view: At early larval stages (∼7 dpf), retinal output has often been conceptualized as functionally homogeneous within a projection class (PC), with each RGC conveying a single dominant response type across its axonal terminals. Middle, updated view: Our analyses reveal that individual RGCs exhibit homogeneous tuning across all terminals, but that multiple RGCs with distinct functional response types (looming, motion, big dot, dimming/OFF, small dot) converge within the same projection class and tectal lamina. Right, later development: By 11 dpf, population-level reorganization continues within this convergent framework. Some response types remain stable, whereas others undergo developmental refinement or reassignment. In particular, big-dot-responsive RGCs can transition toward motion-responsive profiles, contributing to the expansion of motion-sensitive outputs, while experience-dependent tuning biases (e.g., horizontal versus vertical small-dot motion) emerge within specific subtypes.

However, to better inspect the effect of visual deprivation, we further performed both pixel- and cluster-wise Pearson correlation analysis between standard condition (LC1-5) and visual deprivation condition (DRLC1-5) at the late stage (**Fig. 7g**, **h**). Interestingly, we observed minimum or no correlation among LC2 and DRLC2, both of which demonstrated a preference for a small dot movement. This difference stemmed from the response in directionality of stimuli: the normally reared LC2 had a bias for horizontal tuning, whereas DRLC2 exhibited vertical tuning (**Fig. 7i**).

Overall, our results suggest that visual experience does not substantially alter the formation of major functional RGC types, but may instead fine-tune specific response properties. Thus, these results indicate that the functional changes of Satb2-positive RGC axons between 7 and 11 dpf are not attributed to the visual experience itself, but rather to the developmental program that occur during this period.

## Discussion

In this study, we examined how functional responses of Satb2-positive RGCs are organized and maintained during development. We found that functional diversity within tectal layer, or projection class, arises from convergence of distinct RGC types with homogeneous tuning, while population-level reorganization and experience-dependent refinement continue at later larval stages (**Fig. 7j**). Comparison of early (ECs) and late (LCs) functional clusters revealed that three response types, encoding brightening, dimming/OFF, and small-object stimuli, were stable both functionally and anatomically across development. These response types likely represent core visual features that are essential for larval survival at early stages, such as detecting changes in ambient illumination and small moving objects in the environment. In contrast, clusters exhibiting overlapping response profiles retained functional flexibility and continued to mature during late larval stage. Notably, EC6 and EC7 transitioned into LC4 and LC5, respectively, with LC5 becoming the dominant functional population by 11 dpf. These response types, associated with big dot and motion sensitivity, may support behaviors such as prey capture that become increasingly prominent and refined as larvae mature. The expansion of LC5, accounting for over two-thirds of classified responses, likely arises through at least two mechanisms: functional re-tuning of existing RGCs and integration of newly generated RGCs with LC5-like profile. Our longitudinal single-cell analysis supports the former mechanism, revealing that EC5 initially responded to large dots but acquired looming sensitivity over time, becoming part of LC5. Such transition from EC5 to LC5 may be caused by the changes in retinal synaptic inputs, for example from bipolar or amacrine cells, which continue to mature during this period^46^. Similar forms of functional reassignment through recruitment rather than elimination have been described in other visual circuits, including mouse binocular pathways^48^.

The longstanding view in developmental neuroscience is that the retina, unlike cortical visual areas, follows an experience-independent maturation program^25,49^. However, recent work in zebrafish has challenged this view, showing that visual experience can influence retinal circuit structure and function during early larval stage^29,30^. These studies primarily focus on the period before 7 dpf. By examining RGCs beyond this early window, our results extend these findings and indicate that while core functional features of RGCs tend to be shaped by intrinsic programs, experience-dependent refinements can still occur up to 11 dpf. Consistent with this view, cluster identities and overall anatomical distributions of Satb2-positive RGCs remained largely stable even under temporary visual deprivation. This suggests that intrinsic developmental programs, such as changes of synaptic input or neurogenesis, play a dominant role in establishing functional RGC types. Nevertheless, visual experience appears to modulate specific tuning properties. For example, small-dot-responsive EC2/LC2 cells exhibited a preference for horizontally moving stimuli under normal rearing, a feature that shifted toward vertical tuning under dark-rearing condition. This plasticity may reflect adaptation to prey-related motion cues such as paramecia swimming horizontally^50^, usually encountered during the onset of active hunting around 6 dpf. Interestingly, unlike zebrafish, horizontal motion selectivity in mouse retina appears largely insensitive to visual deprivation and instead depends on spontaneous retinal activity^51^. However, the underlying mechanisms in zebrafish retina remain elusive and may need further investigation.

Our study also revises the general view of how functional information is organized within the optic tectum. Although Satb2-positive RGCs belong to defined projection classes and terminate in specific laminae (PC2, PC6, PC8)^4^, we found that multiple functional response types converge within the same tectal layer across development. This organization contrasts with general view of one-to-one correspondence between RGC axonal projection class and functional types^12^. Importantly, single-cell functional imaging analysis revealed that an individual RGC exhibits homogeneous tuning and functional diversity within a layer therefore arises from the convergence of multiple RGC types rather than from mixed tuning within single axons. This single-layer, multi-functional convergence may be critical for the efficient integration of retinal inputs by postsynaptic tectal cells. Zebrafish tectal neurons show diverse dendritic morphologies, with some dendrites arborizing only in one layer (stratified) and others spanning multiple layers of the optic tectum (bushy, non-stratified)^52,53,8,5^. While it is expected that non-stratified tectal dendrites can receive and integrate parallel RGC inputs from multiple layers as recently shown in the mouse^54^, it remains unclear whether stratified tectal neurons receive functionally homogeneous or heterogeneous inputs. Our fundings suggest that even lamina-restricted tectal dendrites may integrate signals from multiple RGC types, potentially broadening the stimulus selectivity of individual tectal neurons. Future studies combining functional imaging and synaptic-resolution anatomy will be needed to explore whether such dendritic computations indeed occur in zebrafish tectal cells.

Finally, our results have implications for understanding the evolutionary conservation and divergence of Satb2-positive RGCs. In mammals, Satb2 marks a restricted set of RGC types, including direction-selective subtype in mouse and rabbit, whereas Satb2-expressing RGCs in primates are morphologically distinct^37^. In zebrafish, we identified that Satb2-positive RGCs comprise a functionally diverse population and do not exhibit strong direction selectivity at early larval stages. However, a subset acquires backward motion preference by 11 dpf, suggesting delayed maturation of direction-related tuning. Unlike canonical direction-selective RGCs, which project to superficial tectal layer^14–16^, Satb2-positive RGC terminate primarily in SFGS3 and SFGS5, indicating that their motion sensitivity may be embedded within broader response profiles rather than forming classical direction-selective circuit. Whether additional direction-selective properties emerge at late developmental stages remain an open question and will require examination of Sab2-positive RGCs in adult zebrafish.

## Acknowledgement

We particularly thank Dr. Dominique Förster for providing custom python scripts. We thank Dr. Tatsumi Hirata, Dr. Takuji Iwasato, Dr. Keisuke Yonehara, Dr. Kazuhide Asakawa, Dr. Yan Zhu, and Dr. Akatsuki Kimura for their discussion and feedback on our manuscript. We thank the members of the Kubo laboratory for their technical assistance and advice as well as the Research Resource Division of RIKEN CBS for fish care and technical assistance. We are grateful to National BioResource Project (NBRP) Zebrafish Japan for providing *Tg(UAS:GFP)* line. AU was supported by the Monbukagaku-sho (MEXT) scholarship. This study was supported by JSPS Grant-in-Aid for Scientific Research (KAKENHI) (17K20147, 21H02586, 22K21353), Naito Foundation and Takeda Science Foundation to FK.

## Author contributions

A.U. and F.K. conceived and designed the study experiments, analyzed the data and developed the project. A.U. performed all the experiments. A.U. and F.K. wrote the manuscript. F.K. supervised and acquired funding for the project.

## Competing interests

Authors declare no competing interests.

## Methods

### CONTACT FOR REAGENT AND RESOURCE SHARING

Further information and requests for resources and reagents should be directed to the Lead Contact Fumi Kubo.

## EXPERIMENTAL MODEL AND SUBJECT DETAILS

### Animal care and transgenic zebrafish

Adult and larval zebrafish (*Danio rerio*) were maintained on a 14 hr light/10 hr dark cycle at 28°C following the standard procedures^55^. All zebrafish experiments were conformed to the guidelines of National Institute of Genetics and RIKEN Center for Brain Science. We used the following previously described transgenic lines: *Tg(UAS:GFP)nns19, Tg(UAS:GCaMP6s)mpn156; Tg(UAS:mCherry)s1984t; Tg(isl2b:GFP)zs7Tg; Tg(elavl3:lyn-tagRFP)mpn404* (a.k.a. *Tg(HuC:lyn-tagRFP)*). For all experiments, embryos were collected and cultured until 6-11 days post fertilization (dpf) in E3 embryo medium supplemented with methylene blue to prevent fungal growth. Sexes of embryos cannot be determined at these developmental stages. For experiments conducted with 11 dpf, larvae were fed ad libitum with paramecia starting from the evening of 7 dpf.

### Generation of *satb2:Gal4* knock-in transgenic zebrafish lines using CRISPR/Cas9 system

Knock-in transgenic zebrafish that express Gal4 gene upstream of candidate genes were generated using previously described protocol^56^. We modified the protocol by using trans-activating CRISPR RNA (tracrRNA, IDT) and synthesized guide CRISPR RNA (crRNA, IDT) instead of short-guide RNA (sgRNA). crRNA1 (for genome digestion), crRNA2 (for donor plasmid digestion), tracrRNA, a donor plasmid and Cas9 protein (IDT) were incubated *in vitro* to from a complex and co-injected into single-cell stage zebrafish embryos. For *satb2*, we designed crRNA1 (TCGTTCTGCTCCAGCGTCTATGG), targeting distinct PAM sequences located within 200-600 base pairs upstream of the transcription site of the *satb2* gene. Concurrent digestion of the genome and the donor plasmid by Cas9 allows the insertion of the donor plasmid containing hsp70 promoter and modified Gal4 gene (Gal4FF) into the promoter region upstream of gene of interest. This integration proceeds via homology independent repair system. For injection we used *Tg(UAS:mCherry)* embryos and pre-screened injected embryos at 3-5 dpf based on mCherry expression in the retina and neuropils in the optic tectum and raised them to adulthood (F0). These potential founder fish were crossed with wild type fish to analyze expression in their offsprings (F1), thus confirming stable transgenesis. Insertions of the Gal4FF transgene into the selected locus were confirmed by PCR. The efficiency for the generation of stable transgenic fish line was 14% (3 out of 21) for *satb2*.

### Sparse labeling of Satb2-positive RGCs

A plasmid DNA *Tol2(UAS:GCaMP6s)* with tol2 mRNA was injected into eggs obtained from *Tg(satb2:Gal4);Tg(UAS:mCherry)* fish at single cell stage. The final concentration constituted 25 ng/μL. The injected embryos were screened at 4-5 dpf using confocal microscope to express both mCherry and GCaMP6s. Samples with minimum number (1 to 3 cells) of GCaMP-positive RGCs were then selected for the functional imaging at 6-7 dpf (see “Two-photon calcium imaging and visual stimulation for functional analysis” below). After completing the functional imaging of sparsely labeled RGCs, whole mount immunohistochemistry was performed to reconstruct the tectal layer arborization pattern of individual RGC.

### Confocal imaging

For live imaging, 6-7 and 10-11 dpf larvae were embedded in 2% low-melting-point agarose and anesthetized in 1% tricaine dissolved in E3 medium. Imaging was performed using confocal microscope (LSM900, Carl Zeiss) with 20x objective. Optical sections were obtained using 0.54-1 µm Z-step.

### Two-photon calcium imaging and visual stimulation for functional analysis

*In vivo* calcium imaging was performed using a two-photon microscope (BergamoII, Thorlabs) equipped with a 20x water-immersion objective (XLUMPLFLN 20x, NA1.0, Olympus) and a 920 nm laser (ALCOR-920, Spark Lasers) on 6-7 and 10-11 dpf *Tg(satb2:Gal4);Tg(UAS:GCaMP6s)* transgenic zebrafish larvae. Larvae were mounted using 2% low-melting-point agarose. Visual stimulus was generated by custom written scripts using PscychoPy3 and projected on white diffusive screen using the red channel of LED projector (DLP LightCrafter 4500) and presented monocularly. Visual stimulation design was adopted and modified from earlier study^5^. Calcium imaging was performed on contralateral optic tectum at 2.187 frames per second at every 10 µm in depth covering most of the tectal neuropil, resulting in typically 8 planes per fish. The pixel size of the imaging was 0.405 µm. Since the average diameter of a presynaptic bouton in zebrafish RGCs is ∼0.8 µm, the physical lateral dimensions of pixels are below that of a typical presynaptic bouton.

Visual stimulation designed by PsychoPy3 consisted of a dark ramp (red to black), a bright ramp (black to red), a dark flash (red to black), and a bright flash (black to red). This was followed by a small horizontally moving dot (5°) presented at five equally spaced elevations across the vertical axis of the screen in forward (temporal to nasal) and backward (nasal to temporal) directions (two repetitions each). Small vertically moving dot (5°) was presented at five equally spaced azimuths across the horizontal axis of the screen in upward (bottom to top) and downward (top to bottom) directions (two repetitions each). Next a big dot was moving horizontally (30°) in forward and backward directions (repeated twice). Dark dots were presented on a bright (red) background for both dot sizes (5° and 30°) moving at a speed of 60°/s. Subsequently, the looming stimuli were presented in two velocities (angular expansion rate of 60°/sec and 20°/sec, corresponding to a fast and a slow loom, respectively) and both ended with a black screen (two repetitions each). Grating stimulus consisting of sinusoidal gratings with a spatial frequency of 0.004 cycle/degree and a temporal frequency of 1.8 Hz moving in 12 equally spaced angular directions was presented in a pseudorandom order. For each presentation of a different direction, the gratings initially stayed stationary for 10 s, in motion for 5 s, and this process was repeated for all grating presentations. The total length of the visual stimulus protocol was about 11 min.

### Pixelwise calcium imaging analysis

Calcium imaging time series was corrected for motion artifacts using TurboReg plugin in ImageJ/FIJI. Neuronal activity of RGC axons in optic tectum was processed using pixelwise analysis based on previous study^5^ using custom written scripts in Python 2.7 and Python 3.2. For each pixel, the fluorescence signal was first normalized to obtain the relative intensity change (ΔF/F₀). 18 regressors representing distinct visual stimulus components were then generated and convolved with a kernel approximating cytosolic GCaMP6s kinetics (τ_decay = 2.5 s). The normalized ΔF/F₀ traces were tested for correlation with each convolved regressor using a linear regression model implemented in the Python scikit-learn package. From the resulting regression coefficients and associated t-statistics, individual t-score maps were generated for each of 18 regressors.

Clustering of responsive RGC pixels was performed using a Python-based pipeline^5^. For each fish, 8 imaging planes (from Z30 to Z100) were included, and pixels with a maximum t-score below 0.6 were excluded. To define general response types, we pooled all responsive pixels (N = 34,350) across six fish imaged at 6-7 dpf and standardized their t-scores based on the 99th percentile per fish. Next, we applied affinity propagation clustering (scikit-learn), with the preference set to the median of pairwise similarities, to the pooled data. The resulting cluster exemplars were then hierarchically organized (using scipy.cluster) with Pearson correlation as the distance metric. Clusters representing less than 3% of all pixels were removed, and a silhouette-informed distance threshold of 0.45 was applied, yielding seven functional early clusters (N = 30,013 pixels). We used the same clustering parameters on data collected at 10-11 dpf and from dark-reared animals at the same age, each yielding five functional clusters.

Unsupervised clustering quality metrics, including the silhouette score and BIC, were used to assess cluster separability within individual datasets obtained from 7 dpf, 11 dpf, and dark-reared 11 dpf larvae (**Extended Data Fig. 1a**, **b**; **Extended Data Fig. 3b**, **c**; **Extended Data Fig. 5b**, **c**). This analysis initially yielded six, five, and five clusters for the 7 dpf, 11 dpf, and dark-reared 11 dpf datasets, respectively. To track cluster reorganization across developmental stages and sensory conditions, we assumed that functional response types are largely stable. To maximize correspondence of clusters across stages, we sought to fix clustering parameters globally, including the dendrogram distance threshold and pixel exclusion criteria. At 7 dpf, the initial metrics favored a six-cluster solution. However, enforcing a six-cluster solution required merging two response profiles that later emerged as clearly separable clusters at 11 dpf and under dark-rearing conditions, thereby disrupting the continuity of cluster identities across development. Under these constraints, a seven-cluster solution at 7 dpf preserved functional identities that could be consistently traced across later stages, supporting its biological validity despite a modest underestimation of the single-dataset clustering metrics. Based on these considerations, we set the dendrogram distance threshold (max_d) to 0.45 and the pixel exclusion criterion to the top 3% of pixels. This approach ultimately yielded seven, five, and five clusters for the 7 dpf, 11 dpf, and dark-reared 11 dpf datasets, respectively.

The seven early clusters (ECs) were used as a reference to classify RGC pixels in individual fish by correlating each pixel’s t-scores to the exemplar profiles. Each pixel was assigned to the best-matching cluster. These assignments were used to visualize spatial distributions and to generate anatomical cluster maps. For inter-fish comparisons, pixel-level cluster maps were registered to a zebrafish standard brain using ANTs-based nonlinear transformation (see “Image registration of functional imaging for anatomical mapping” section). The same pipeline was applied to functional imaging data collected at 11 dpf, yielding five late clusters (LCs), and was also used to identify five LCs from dark-reared fish (DRLC) of the same age.

To assess single-RGC functional profiles, responses from sparsely labeled axons were first analyzed using the similar pixel-wise analysis using the EC template derived at 7 dpf. For longitudinal analysis, the same individual RGCs imaged at both 6-7 dpf and 10-11 dpf were analyzed using the 7 dpf EC template for early timepoints and the 11 dpf LC template for later timepoints, respectively. This enabled the direct comparison of cluster identity and functional maturation at single-cell resolution.

To assess the stability and similarity of RGC response clusters across developmental stages, two complementary analyses were performed: pixel-level similarity and inter-cluster similarity. For pixel-level similarity, a comprehensive correlation matrix was generated by computing the Pearson correlation coefficient (r) between all individual pixels pooled from both time points (7 and 11 dpf). Pairwise correlations were calculated based on each pixel’s responses to 18 regressor. The resulting heatmap shows high within-cluster similarity along the diagonal, and varying degrees of inter-cluster similarity in off-diagonal regions. Pixels were ordered by cluster assignment. For inter-cluster similarity, the average Pearson correlation coefficient (r) was computed between every pair of clusters across same or different developmental time points. This was done by comparing the response scores to 18 stimulus regressors, averaged across all pixel pairs between each cluster pair. The resulting similarity matrix reveals the degree of correspondence between early and late cluster response profiles, with r-values ranging from -1 to 1, where r = 1 indicates perfect positive correlation, r = -1 indicates perfect negative correlation, and r = 0 indicates no correlation.

### Image registration of functional imaging for anatomical mapping

After calcium imaging, anatomical Z-stacks of the same fish *Tg(satb2:Gal4);Tg(UAS:GCaMP6s);Tg(HuC:lyn-tagRFP)* were acquired. First, small stacks (256×256 pixels, Z-step size: 0.57-1 μm) were collected using a two-photon microscope to capture the GCaMP6s (green) channel. Subsequently, an overview stack was acquired using a confocal microscope with a 20× objective (512×512 pixels, Z-step size: 0.57-1 μm), capturing both GCaMP6s (green) and lyn-tagRFP (red) channels. To visualize and compare functional activity across multiple fish in a common reference space, we implemented a three-step image registration pipeline^14^: 1) Functional-to-anatomical registration: The functional two-photon Z-stacks were first coarsely aligned to the corresponding confocal anatomical stacks using manual landmark selection (Fiji plugin “Name Landmarks and Register”; Longair & Jefferis), using 5-6 reference points. Fine registration of the two-photon GCaMP6s stack to the confocal GCaMP6s channel was then performed using the Advanced Normalization Tools (ANTs) library^57^ (version 2.6.3), with parameters reported in previous study^58^ and the antsRegistration.sh command. Functional data, representing cluster-specific pixel activity at 8 Z-planes (seven clusters for 7 dpf, five clusters for 11 dpf), were binned into template Z-stacks matching the anatomical dimensions using custom Python script. These were subsequently aligned to the confocal stack using the antsApplyTransforms.sh command. 2) Anatomical-to-standard registration: The confocal anatomical stacks were registered to a standard live zebrafish brain *Tg(HuC:lyn-tagRFP)* at 6 dpf obtained from mapzebrain.org, using the lyn-tagRFP (red) channel as a reference and the antsRegistration.sh command for alignment. For 11 dpf data, standard brain was generated from six different stacks of *Tg(HuC:lyn-tagRFP)* at 10 dpf using the command antsMultivariateTemplateConstruction2 in ANTs. 3) Functional-to-standard registration: Finally, the anatomical-to-standard transformation parameters were applied to the registered functional stacks using the antsApplyTransforms.sh command, thus mapping functional pixel data from each fish into the common reference brain space. This pipeline enabled consistent registration of functional signals from all six fish (N = 6) to a standardized anatomical framework. For subsequent 3D rendering, we used the open-source software 3D Slicer^59^ (version 5.6.2) to load the affine-registered functional pixels from all fish and the standard brain. The corresponding color codes and opacity were manually assigned for visualization in accordance with each cluster identity.

### Anteroposterior (AP) and dorsoventral (DV) distribution analysis

To quantify the spatial distribution of Satb2-positive RGC terminals along the anteroposterior (AP) and dorsoventral (DV) axes of the tectum, we performed the following analyses. For each fish, the tectal neuropil was manually delineated in every imaging plane to generate a binary mask that defined the anatomical boundaries. For anteroposterior (AP) axis computation, the principal AP axis was estimated by applying singular value decomposition (SVD) to the spatial coordinates of pixels within the mask. The first eigenvector defined the AP axis, and the centroid of the mask served as the origin. Functionally classified pixels were projected onto this AP axis, and their positions were normalized to a range of -0.5 (posterior) to +0.5 (anterior). Distributions of pixel positions were visualized by kernel density estimation (KDE), weighted by the relative fraction of pixels belonging to each cluster. For each cluster, the mode of the KDE (i.e. highest estimated density) was taken as the dominant position along the AP axis. Kernel density estimation (KDE) rather than histograms was used to avoid artifacts introduced by arbitrary binning and to obtain smooth, continuous distributions of pixel density.

In case of dorsoventral (DV) axis computation, the DV axis was defined according to imaging depth, using registered Z-planes spanning from dorsal (Z30-Z40) to ventral (Z90-Z100) tectum. Each Z-plane was assigned a normalized DV coordinate using a discrete binning scheme (0.0-0.7), in which each plane was assigned a fixed DV index (0 = ventral-most, 7 = dorsal-most). Each functionally classified pixel inherited the DV coordinate of its parent Z-plane. As with AP, cluster-wise KDEs were computed across all pixels, and the KDE mode was used to indicate the dominant DV position for each cluster. Pixel densities were normalized by the total number of pixels across clusters to enable direct comparison of relative distributions.

### Quantification of Direction and Orientation Selectivity

To assess the visual tuning properties of Satb2-positive RGCs, calcium imaging data were analyzed pixel-wise at 7 and 11 dpf across 6 fish per time point. For each pixel within Satb2-positive regions, responses to 12 drifting grating stimuli presented in different directions were extracted. Direction Selectivity Index (DSI) and Orientation Selectivity Index (OSI) were calculated as the normalized vector sum of responses across directions or orientations, respectively, as previously described with some modifications^14^. Pixels with DSI or OSI values exceeding 0.5 were classified as direction- or orientation-selective. For direction-selective pixels, the preferred direction (0°-360°) was identified as the angle of the peak response, while for orientation-selective pixels, the preferred orientation (0°-180°) was determined. Histograms were generated to visualize the distribution of selectivity indices and preferred directions/orientations. Population-level tuning was summarized using polar plots, where each radial bin indicates the number of selective pixels tuned to a given direction or orientation.

### Whole mount immunohistochemistry (IHC)

The IHC staining protocol for whole mount samples was adopted from previous study^60^. In short, after 2-photon calcium imaging zebrafish larvae were fixed overnight in a solution of 4% paraformaldehyde in PBS at 4°C. For the antigen retrieval the following day, samples were washed in PBT (PBS with 0.25% TritonX), incubated in 150 mM Tris-HCl pH 9 for 5 minutes at room temperature, and then transferred to 63°C for 15 minutes. After the wash with PBT, samples were incubated in cold Trypsin-EDTA on ice for 45 minutes and washed again. For the blocking step, samples were immersed into the blocking solution (PBT with 1% DMSO, 1% BSA and 5% Serum) for 3 hours at room temperature. Then, whole mount larvae were first incubated in the primary antibody solution (chicken anti-GFP (A10262, Thermo Fisher Scientific/Invitrogen, 1:1000), mouse DsRed (632393, Clontech/Takara, 1:500), rabbit anti-Synapsin 1/2 (106002, Synaptic Systems, 1/1000)) for 7 days at 4°C, followed by the secondary antibody solution (goat anti-chicken Alexa Fluor 488 (A11039, Thermo Fisher Scientific/Invitrogen, 1:250), goat anti-mouse Alexa Fluor 555 (A21422, Thermo Fisher Scientific/Invitrogen, 1:250), goat anti-rabbit Alexa Fluor 647 (A21244, Thermo Fisher Scientific/Invitrogen, 1:250), DAPI (1:1000)) for 5 more days at 4°C. After washing and post-fixing in 4% PFA, samples were maintained in 87% glycerol at 4°C.

### Hybridization Chain Reaction FISH staining

Hybridization Chain Reaction (HCR) RNA-FISH was performed following a modified version of the Molecular Instruments protocol described previously^61^. 6-7 dpf larvae were fixed in 4% PFA overnight at 4°C, washed in DPBST, and briefly treated with cold methanol for permeabilization. After rehydration, samples were prehybridized, then incubated with HCR Satb2 probe solution (2 pmol per 500 µl hybridization buffer) for 12-16 hours at 37°C. Following stringent washes, amplification was carried out using snap-cooled hairpins (5 pmol h1 and h2 per 250 µl amplification buffer) overnight at room temperature in the dark. After final washes, larvae were stored in 5× SSCT at 4°C until imaging.

## Data and code availability

Data and code will be made available upon request.

## Extended data

**Extended Data Figure 1.**
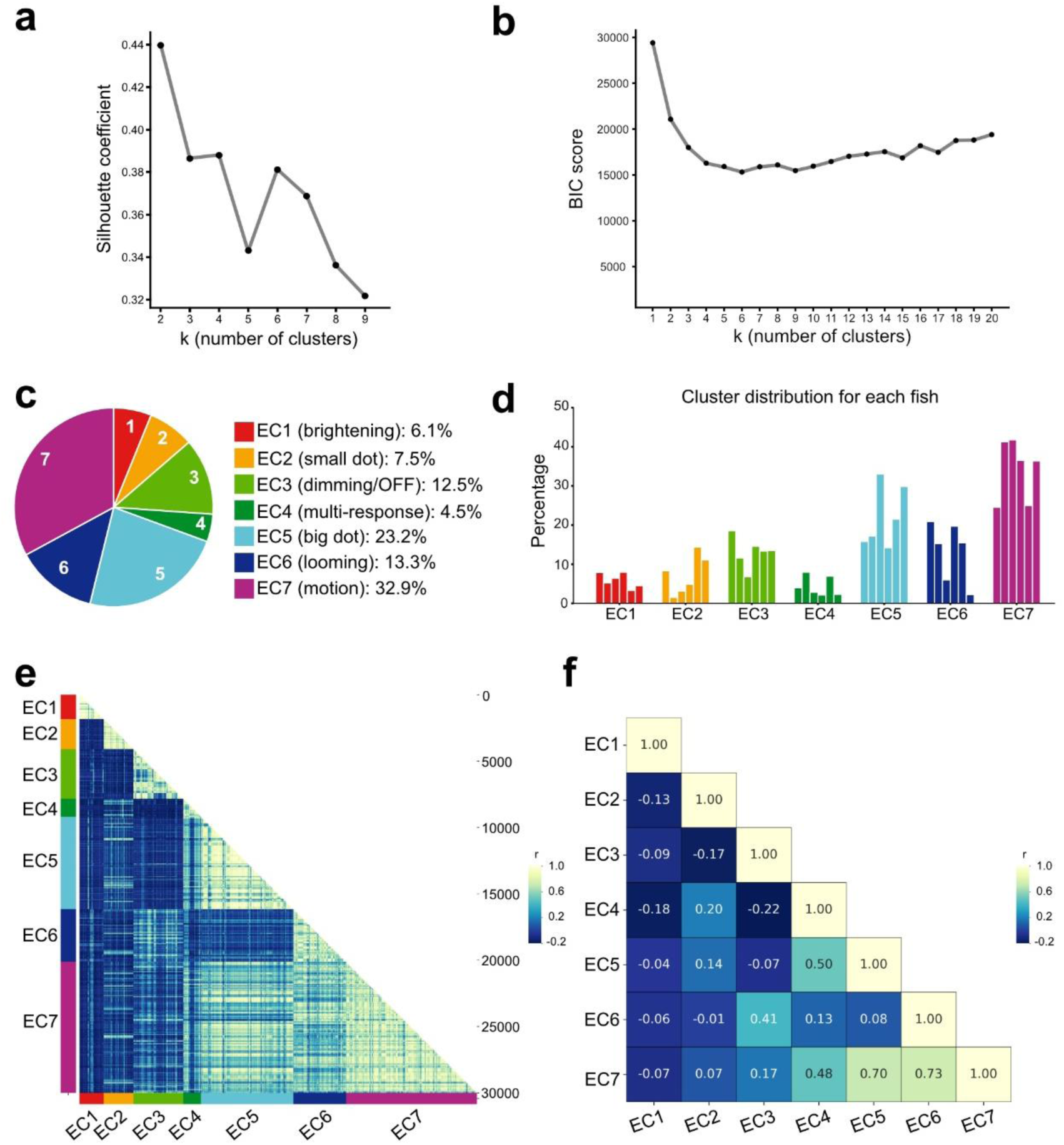
Functional characterization of Satb2-positive RGCs using clustering analysis. **a)** Silhouette plot for cluster validation. The silhouette score shows a secondary local maximum at k = 6, indicating improved separation at this cluster number at 7 dpf. **b)** Bayesian Information Criterion (BIC) evaluated across different numbers of clusters from 7 dpf. BIC values reach a minimum around k = 6, beyond which additional clusters provide diminishing improvements in model fit. **c)** Pie chart showing proportion of all selected pixels from functional clusters with representative response type and percentage indicated along the corresponding Early Cluster (EC1-7) from 7 dpf. **d)** Bar chart showing distribution of functional Satb2-positive RGC pixels corresponding to color-coded EC1-7 among 6 fish obtained at 7dpf. **e)** Pixel-to-pixel correlation heatmap displaying Pearson correlation coefficient (r) between all individual pixel cluster of each EC from 7 dpf. Pixels are ordered by cluster assignment in Fig.2d. For each cluster pixels group, the mean Pearson correlation was calculated using their response scores to 18 stimulus regressors. **f)** Cluster-to-cluster similarity matrix based on the Pearson correlation coefficient (r) between each pair of ECs identified at 7 dpf.

**Extended Data Figure 2.**
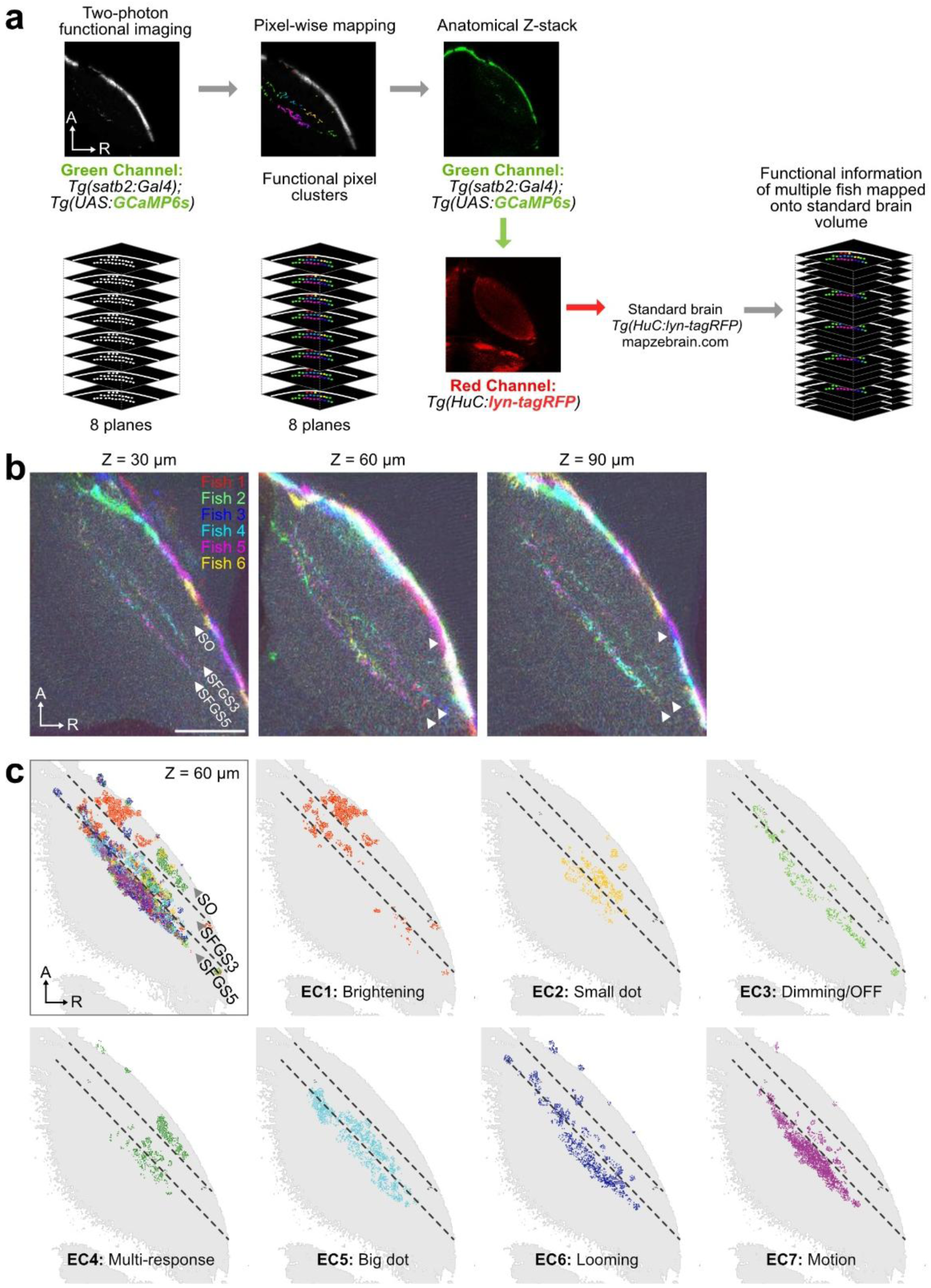
Anatomical characterization of Satb2-positive RGCs using clustering analysis. **a)** Workflow for mapping functionally classified Satb2-positive RGC responses to a common reference brain. Left: Two-photon calcium imaging was performed in Satb2-expressing RGC terminals (green: *Tg(satb2:Gal4);Tg(UAS:GCaMP6s)* in the optic tectum across 8 imaging planes. Middle: Pixel-wise mapping was used to assign individual pixels to functional clusters based on their visual response profiles. Right: Functional imaging planes were registered to a high-resolution anatomical Z-stack from the same fish, and subsequently aligned to a standard zebrafish brain volume *Tg(HuC:lyn-tagRFP)* from mapZebrain atlas. This pipeline enables integrated visualization and comparison of functional RGC responses across multiple fish within a unified anatomical framework. **b)** Cross-sectional images (Z-slices) at three depths (Z = 30 μm, 60 μm, 90 μm) showing registered brains of six individual *Tg(satb2:Gal4);Tg(UAS:GCaMP6s)* fish, with functionally classified Satb2-positive RGC pixel clusters mapped into a common reference brain (*Tg(HuC:lyn-tagRFP)* from mapZebrain). Each fish is color-coded (Fish 1: red, Fish 2: blue, Fish 3: cyan, Fish 4: green, Fish 5: magenta, Fish 6: yellow). The spatial overlap of RGC axons across fish demonstrates robust anatomical registration. Z-position values indicate depth from the dorsal-most surface. A, anterior; R, right. Scale bar, 50 µm. **c)** Spatial distribution of individual early response clusters within tectal laminae at Z = 60 μm depth. Each cluster is shown separately with corresponding functional annotation (EC1-7). A, anterior; R, right.

**Extended Data Figure 3.**
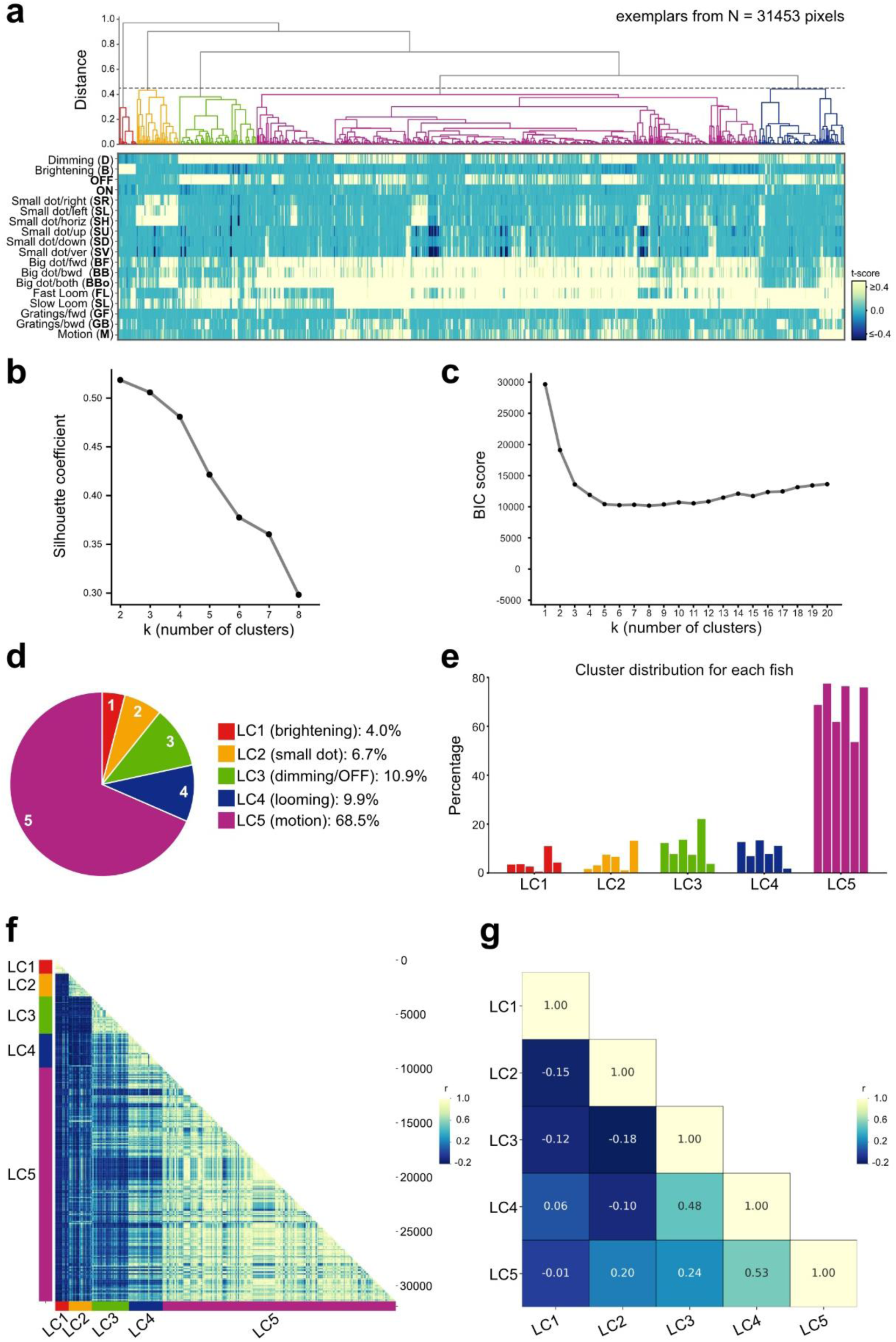
Functional and anatomical characterization of Satb2-positive RGCs over development. **a)** Hierarchical clustering dendrogram and corresponding heatmap of response profiles from N = 31,453 pixels, aligned by regressor responses of functional Satb2-positive RGC pixels in the optic tectum at 11 dpf. Each column represents a single pixel’s activity across stimuli and each row is a stimulus condition (listed on the left). A grey dotted line indicates a chosen distance threshold of 0.45, which yielded five functional clusters named as Late Cluster (LC1-5) with a distinct color code. The heatmap shows responses of each pixel to 18 regressor types listed and abbreviated along y axis. **b)** Silhouette plot for cluster validation. The silhouette score shows a chosen maximum at k = 5 at 11 dpf. **c)** Bayesian Information Criterion (BIC) evaluated across different numbers of clusters from 11 dpf. BIC values reach a minimum around k = 5. **d)** Pie chart summarizing the proportion of pixels belonging to each functional LC from 11 dpf. **e)** Bar chart showing distribution of functional Satb2-positive RGC pixels corresponding to color-coded LC1-5 same as in (a) among 6 fish obtained at 11 dpf. **f)** Pixel-to-pixel correlation heatmap displaying Pearson correlation coefficient (r) between all individual pixel cluster of each LC from 11 dpf. Pixels are ordered by cluster assignment in (a). For each cluster pixels group, the mean Pearson correlation was calculated using their response scores to 18 stimulus regressors. **g)** Cluster-to-cluster similarity matrix based on the Pearson correlation coefficient (r) between each pair of LCs identified at 11 dpf.

**Extended Data Figure 4.**
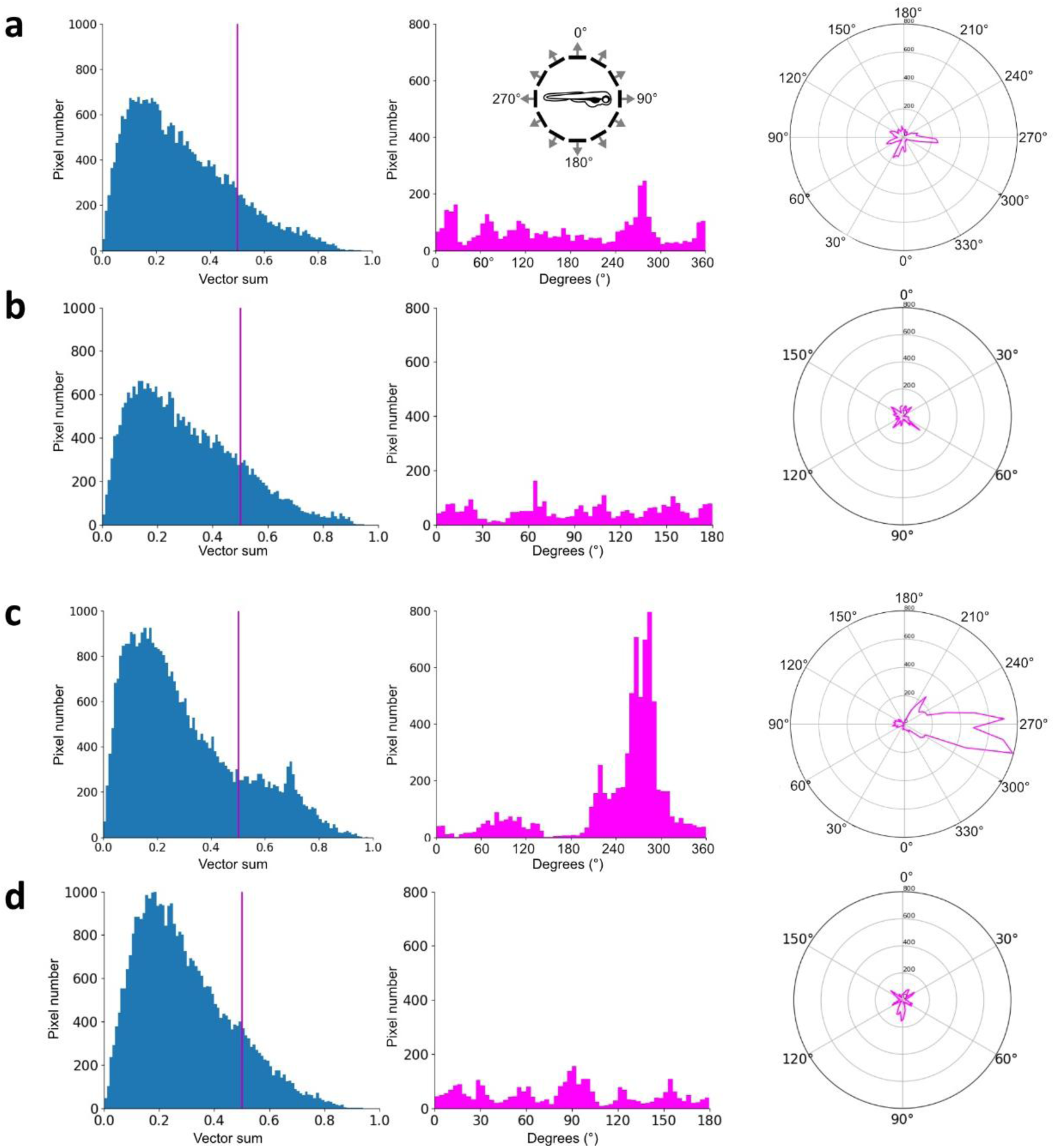
Developmental changes in direction and orientation tuning in Satb2-positive RGCs. **a)** Distribution of direction-selective responses in Satb2-positive RGC pixels at 7 dpf (N = 6 fish). The left histogram shows the vector-sum magnitude for each pixel across the 12 drifting grating directions; a magenta line marks the threshold (direction selectivity index, DSI > 0.5) used to classify pixels as direction-selective. The middle panel shows a histogram of the preferred motion directions (0°-360°) of pixels exceeding this threshold. The right panel is a polar plot summarizing the population tuning: each radial bin corresponds to a motion direction, and its length indicates the number of pixels preferring that direction at 7 dpf. **b)** Orientation-selective responses of Satb2-positive RGCs at 7 dpf (N = 6 fish). The left histogram depicts the vector-sum magnitude for each pixel and magenta line marks the threshold (orientation selectivity index, OSI > 0.5) used to classify pixels as orientation-selective. The middle panel shows a histogram of preferred orientations (0°-180°) among pixels above this threshold. The right panel is a polar plot summarizing the population orientation preference and each radial axis corresponds to an orientation and its length reflects the number of pixels tuned to that orientation at 7 dpf. **c)** Direction-selective responses of Satb2-positive RGCs at 11 dpf (N = 6 fish). This mirrors the analysis in Panel (a). **d)** Orientation-selective responses of Satb2-positive RGCs at 11 dpf (N = 6 fish). This mirrors the analysis in Panel (b).

**Extended Data Figure 5.**
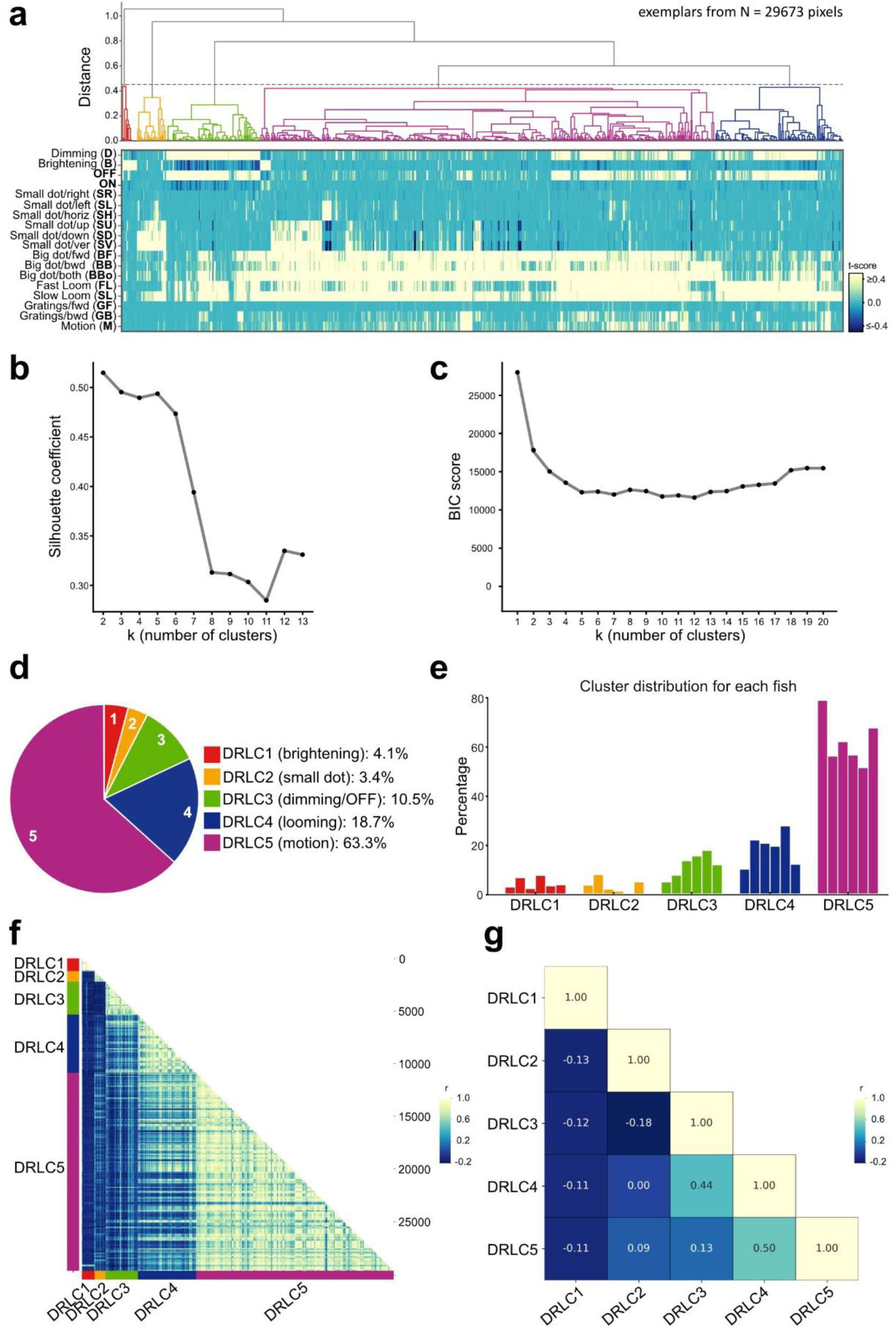
Functional and anatomical characterization of Satb2-positive RGCs over development under dark-reared condition. **a)** Hierarchical clustering dendrogram and corresponding heatmap of response profiles from N = 29,673 pixels, aligned by regressor responses of functional Satb2-positive RGC pixels in the optic tectum at 11 dpf under dark-reared condition. Each column represents a single pixel’s activity across stimuli and each row is a stimulus condition (listed on the left). A grey dotted line indicates a chosen distance threshold of 0.45, which yielded five functional clusters named as Dark-Reared Late Clusters (DRLC1-5) with a distinct color code. The heatmap shows responses of each pixel to 18 regressor types listed and abbreviated along y axis. **b)** Silhouette plot for cluster validation. The silhouette score shows a chosen maximum at k = 5 at 11 dpf under dark-reared condition. **c)** Bayesian Information Criterion (BIC) evaluated across different numbers of clusters from 11 dpf under dark-reared condition. BIC values reach a minimum around k = 5. **d)** Pie chart summarizing the proportion of pixels belonging to each functional DRLCs from 11 dpf. **e)** Bar chart showing distribution of functional Satb2-positive RGC pixels corresponding to color-coded DRLC1-5 same as in (a) among 6 fish obtained at 11 dpf. **f)** Pixel-to-pixel correlation heatmap displaying Pearson correlation coefficient (r) between all individual pixel cluster of each DRLCs from 11 dpf. Pixels are ordered by cluster assignment in (a). For each cluster pixels group, the mean Pearson correlation was calculated using their response scores to 18 stimulus regressors. **g)** Cluster-to-cluster similarity matrix based on the Pearson correlation coefficient (r) between each pair of DRLCs identified at 11 dpf.

## Notes

### Competing Interest Statement

The authors have declared no competing interest.

